# Targeting TACC3 represents a novel vulnerability in highly aggressive breast cancers with centrosome amplification

**DOI:** 10.1101/2022.05.18.492567

**Authors:** Ozge Saatci, Ozge Akbulut, Metin Cetin, Vitali Sikirzhytski, Ozgur Sahin

## Abstract

Centrosome amplification (CA) is a hallmark of cancer that is strongly associated with highly aggressive disease and worse clinical outcome. However, there are no effective strategies targeting cancer cells with CA while sparing normal cells. Here, we identified Transforming Acidic Coiled-Coil Containing Protein 3 (TACC3) to be overexpressed in tumors with CA, and its high expression is associated with dramatically worse clinical outcome. We demonstrated that TACC3 forms distinct functional interactions in mitotic and non-mitotic cancer cells with CA to facilitate centrosome clustering (CC) and transcriptional repression of tumor suppressors, respectively. We showed, for the first time, that TACC3 interacts with the Kinesin Family Member C1 (KIFC1) via its TACC domain in mitotic cells with CA and inhibition of TACC3 blocks this interaction, leading to apoptosis via multipolar spindle formation and activation of spindle assembly checkpoint (SAC)/CDK1/p-Bcl2 axis. In interphase, TACC3 interacts with the members of the nucleosome remodeling and deacetylase (NuRD) complex (HDAC2 and MBD2) in nucleus, and its inhibition causes p53-independent G1 arrest and apoptosis by blocking these interactions and activating the transcription of key tumor suppressors (e.g., p21, p16 and APAF1). Notably, inducing CA by chemical (cytochalasin D) or genomic (PLK4 overexpression or p53 loss) modulations renders cancer cells highly sensitive to TACC3 inhibition. Targeting TACC3 by small molecule inhibitors or guide RNAs strongly inhibits growth of organoids and breast cancer cell line- and patient-derived xenografts with CA. Altogether our results pave the way towards therapeutic targeting of TACC3 in highly aggressive cancers.

## INTRODUCTION

Centrosome amplification (CA) is highly prevalent in cancer^1,2^. It is strongly associated with tumor progression and worse prognosis in a variety of different cancers, e.g., breast, prostate, ovarian and lung^2^. Preclinical studies demonstrated that supernumerary centrosomes can trigger tumor initiation^3^ as well as cancer cell invasion^4^. There are various causes of CA, including centrosome overduplication by PLK4 overexpression, cytokinesis failure or genomic aberrations, such as p53 mutations^5,6^ that are commonly observed in aggressive tumors, e.g., triple negative breast cancer (TNBC)^7^. Given the high prevalence of CA and its association with tumor aggressiveness, therapeutics targeting cancer cells with supernumerary centrosomes (described as “Achilles’ heel of cancer”^8^) represent a unique opportunity to eliminate the most aggressive tumors while sparing the normal cells.

Centrosomes are the major microtubule organizing centers within the cells. During mitosis, centrosomes aid in formation and orientation of the mitotic spindles and support bipolar division. Clustering extra centrosomes (CC) into opposite spindle poles during mitosis is critical for preventing multipolar division and apoptosis in cancer cells with CA^9^. Members of the chromosomal passenger complex (CPC), e.g., Aurora A kinase, the kinetochore component, Ndc80 complex, and the minus end-directed motor protein, KIFC1 have been previously shown to facilitate CC and mitotic progression in cancer cells with CA^9,10^. However, in clinical settings, such mitosis-directed strategies have been unsuccessful with poor efficacy, potentially due to low fraction of mitotic cells within the tumors^11^. Furthermore, severe side effects associated with targeting of these proteins due to their multifaceted functions in several diverse processes^12,13^ hinder their clinical development. Therefore, identifying and targeting novel proteins essential for both mitotic and non-mitotic cancer cells are necessary that will interfere with not only mitosis but also disrupt interphase-related processes to achieve long-lasting anti-tumorigenic effect with minimal toxicity^11^.

Transforming acidic coiled-coil 3 (TACC3) is upregulated in solid tumors and strongly associated with worse prognosis in several different cancers, such as breast^14^, lung^15^ and ovarian cancer^16^. It is localized to centrosomes as well as microtubules and controls spindle stability and microtubule nucleation^17,18^. Inhibiting TACC3 in *in vitro* settings has been shown to cause spindle defects and mitotic catastrophe^17^. We recently reported a potent TACC3 inhibitor (BO-264) that impaired tumor growth in mouse tumor models and increased survival without any major toxicity^19^. Despite the growing body of evidence reporting the prognostic power of TACC3 in cancer, the therapeutic potential of targeting TACC3 to inhibit the growth of highly aggressive tumors with CA has not been studied. Furthermore, little is known if TACC3 has critical functions beyond mitosis via interacting with distinct functional partners in different cell cycle stages.

In this study, we identified TACC3 as being strongly upregulated in cancers with CA and associated with worse clinical outcome in highly aggressive patient subpopulations. We demonstrated a strong CA-directed vulnerability to TACC3 inhibition in organoids, cell line- and patient-derived xenografts with CA using genomic knockout and pharmacologic inhibition of TACC3. We showed that TACC3 drives tumorigenesis by forming distinct functional interactions in different phases of cell cycle. It interacts with KIFC1 in mitotic CA cells, mediating CC; while in interphase, it interacts with the members of the nucleosome remodeling and deacetylase (NuRD) complex to suppress the transcription of key tumor suppressors, promoting G1/S transition and inhibiting apoptosis. Therefore, targeting TACC3 inhibits these interactions and on one hand, causes centrosome de-clustering and mitotic catastrophe via activating the SAC/CDK1/p-Bcl2 axis, and on the other hand, induces p53-independent G1 arrest, ultimately causing apoptotic cell death.

## RESULTS

### High TACC3 correlates with disease aggressiveness in patients with CA and breast cancer cells with CA are highly sensitive to TACC3 inhibition

To analyze the expression of TACC3 in breast cancer patients with CA, patients were stratified based on their CA status by using a published gene signature of CA (CA20). This list contains 20 genes that have been experimentally demonstrated to cause CA when dysregulated^20^. Importantly, TACC3 is not in this gene signature. As shown in **Fig. 1A**, TACC3 is higher in breast cancer tumors with high CA20 score in the METABRIC dataset^21^, and among high CA patients, those who express higher TACC3 exhibit much worse survival as compared to those who express lower TACC3 (**Fig. 1B**), suggesting that TACC3 may be critical for the outcome of patients with CA. Importantly, CA20 score and TACC3 levels were higher in HER2+ and basal subtypes of breast cancer which are the two most aggressive subtypes (**Fig. 1C, D**). These results were further validated in an independent breast cancer dataset, GSE25066 (**fig. S1A-D**). In a panel of breast cancer cells, we observed that cancer cell lines with CA (mostly TNBC and HER2+) express higher levels of TACC3 and are more sensitive to TACC3 inhibition with BO-264^19^ (**Fig. 1E-G**). Furthermore, a TNBC PDX organoid expressing TACC3 and bearing CA (**Fig. 1H**) responded to BO-264 with a nanomolar range IC50 (**Fig. 1I**). Furthermore, TACC3 expression significantly correlates with CA20 score in the pan-cancer CCLE dataset, comprising the gene expression profiling of more than 1000 cell lines from 36 different cancer types (**fig. S1E**). Importantly, we showed that TACC3 is higher (*P*<0.001) and it is associated with worse survival in CA tumors of not only breast cancer but also of prostate (*P*=0.001, HR=7.25), lung (*P*=0.049, HR=2.19) and head & neck (*P*=0.13, HR=1.96) cancers, suggesting that its association with CA is more general (**fig. S1F-K**).

**Figure 1.**
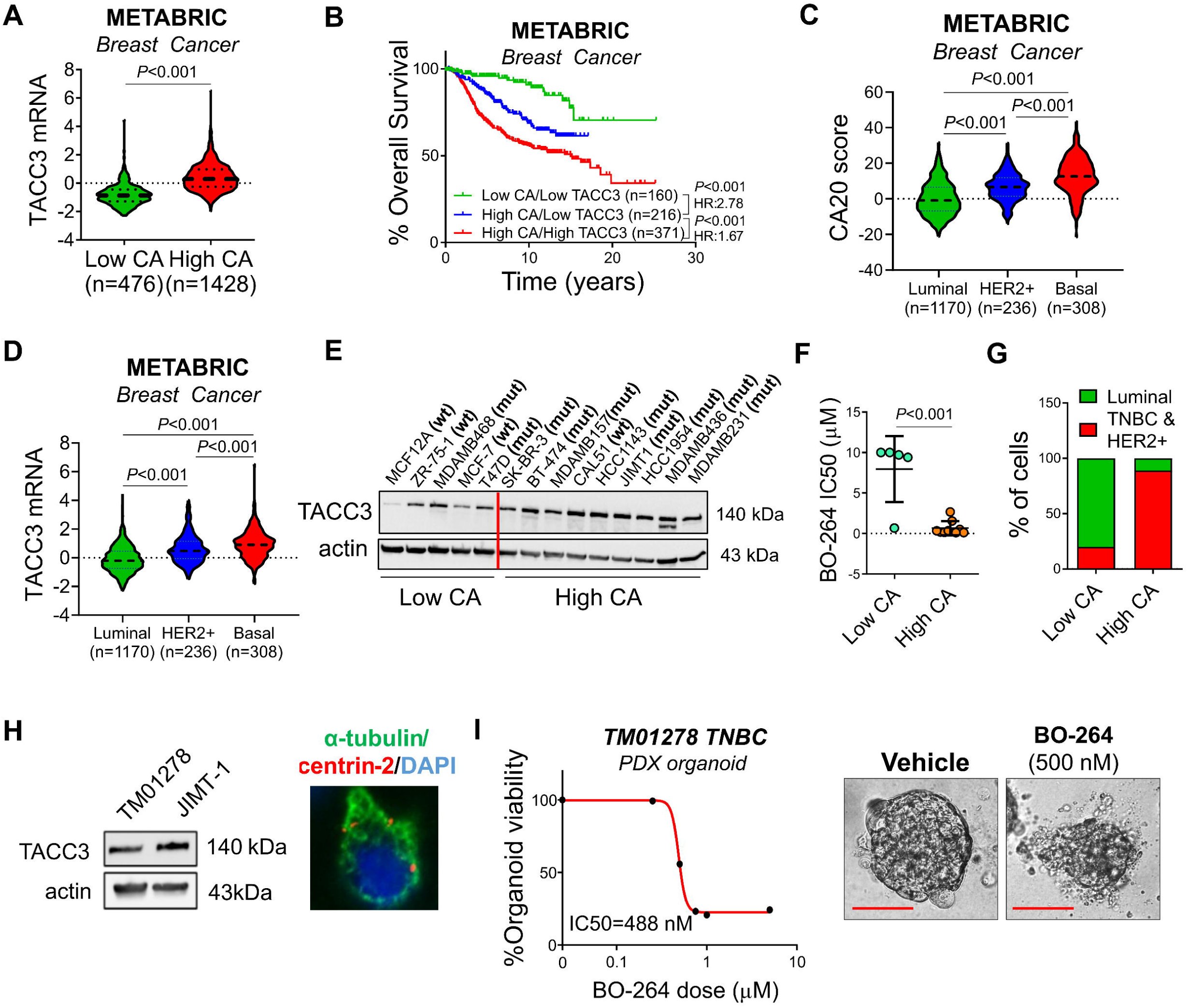
TACC3 correlates with disease aggressiveness in patients with CA and breast cancer cells/organoids with CA are highly sensitive to TACC3 inhibition. (**A**) TACC3 mRNA expression in breast cancer tumors in the METABRIC dataset with low vs. high CA20 score. (**B**) Survival analyses in breast cancer patients based on CA score and TACC3 expression. (**C, D**) Expression of CA20 score (C) and TACC3 mRNA (D) in different breast cancer subtypes in METABRIC dataset. (**E**) TACC3 expression in a panel of breast cancer cell lines with CA and no CA. (**F**) Sensitivity of breast cancer cell lines from E to TACC3 inhibitor, BO-264 with respect to CA status. (**G**) Percentage of breast cancer cell lines from E belonging to TNBC and HER2+ subtype vs. luminal subtype based on CA status. (**H**) Western blot analysis of TACC3 in TNBC PDX TM01278 organoids in comparison with JIMT-1 cells (left panel) and α-tubulin (green) and centrin 2 (red) staining in TM01278 organoids showing CA (right panel). (**I**) Dose response curve of TM01278 organoids upon BO-264 treatment for 1 week. The representative images are provided on the right panel. The scale bar is 100 µm. CA, centrosome amplification; HR, hazard ratio.

### TACC3 is associated with centrosome clustering (CC) in patients with CA and mediates CC in cells with CA

Centrosome clustering (CC) is a critical process required for bipolar division and survival of cancer cells with CA. To test if TACC3 has a role in CC, we examined the correlation of TACC3 with a CC-related gene signature^9^ in breast cancer patients with CA. We observed that patients with CA that have high TACC3 expression, and therefore have worse survival (as shown in **Fig. 1B**) exhibit higher levels of the CC score (**Fig. 2A**). These results were also validated in other tumors with CA, i.e., prostate, lung, and head & neck cancers (**fig. S2A-C**). Notably, TACC3 expression correlates with CC score in the pan-cancer CCLE dataset (**fig. S2D**) and the NCI60 cell line panel (**fig. S2E**). Cancer cells having high CC scores from the NCI60 panel were shown to be more sensitive to TACC3 inhibitor, BO-264 (**fig. S2F**), further suggesting the critical involvement of TACC3 in mediating CC and cell survival.

**Figure 2.**
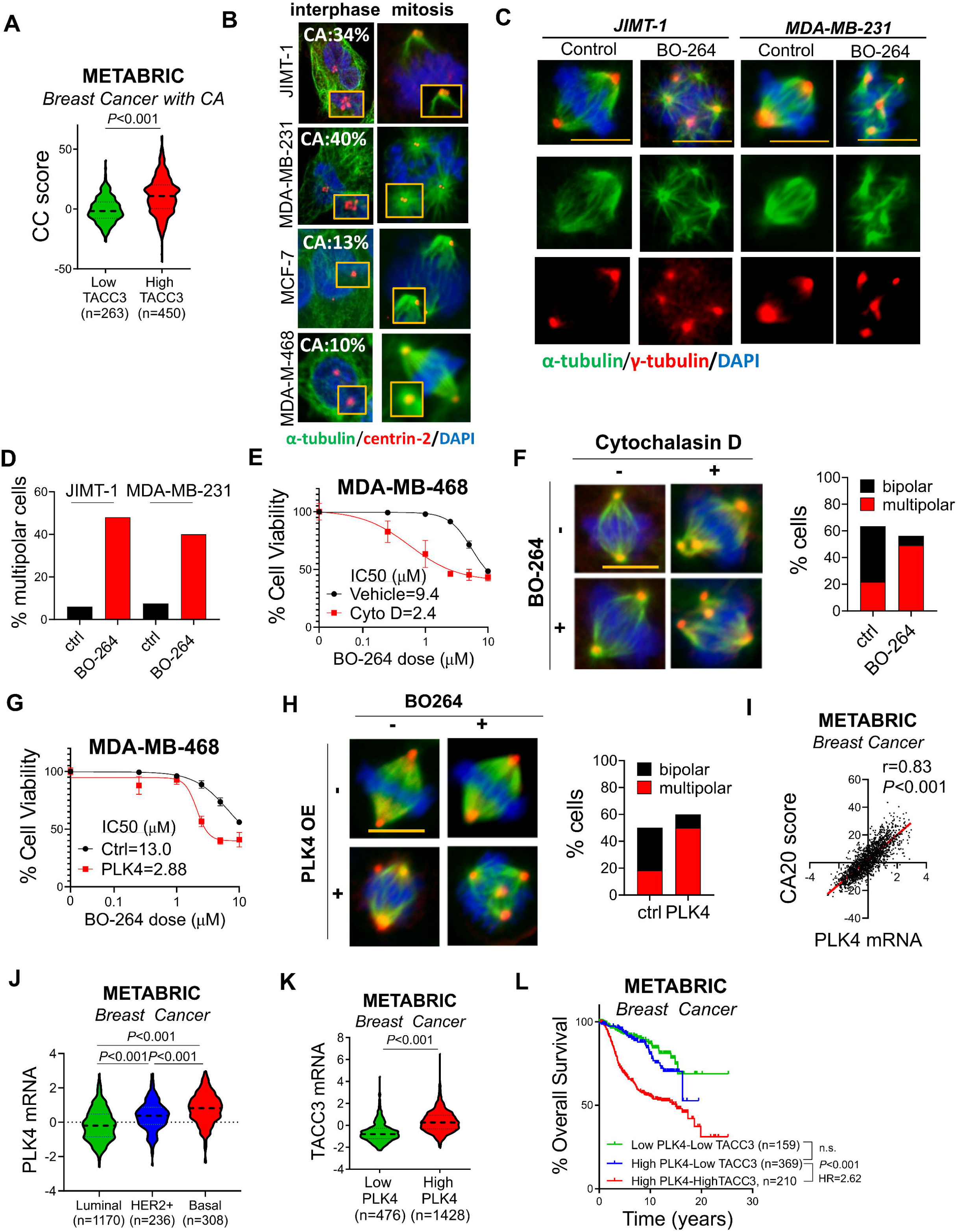
TACC3 correlates with centrosome clustering (CC) in patients with CA and mediates CC in cells with CA. (**A**) Expression of CC score in breast tumors with CA from METABRIC with low vs. high TACC3 expression. (**B**) CA status (as % of cell population) of breast cancer cell lines determined by centrin-2 (red) and α-tubulin (green) in interphase and mitosis. (**C**) Multipolar spindle formation in BO-264-treated JIMT-1 and MDA-MB-231 cells as shown by α-(spindle, green) and γ-(centrosome, red) tubulin staining. Scale bar=10 µm. (**D**) Quantification of mitotic cells with multipolar spindles from C. (**E**) Dose response curve of MDA-MB-468 cells 72 hrs after treatment with BO-264 upon CA induction with 1 µM of cytochalasin D for 20 hrs. (**F**) IF staining of α-(green) and γ-(red) tubulin in MDA-MB-468 cells treated with 1 µM of cytochalasin D for 20 hrs followed by 24 hrs treatment with 5 µM BO-264. Scale bar=10 µm. Quantification of mitotic cells with multipolar spindles is provided on the right panel. (**G**) BO-264 dose response curve of MDA-MB-468 cells transfected with control vector or PLK4 to induce CA. (**H**) IF staining of α-(green) and γ-(red) tubulin in MDA-MB-468 cells transfected with control or PLK4 vector for 24 hrs followed by treatment with 5 µM BO-264 for an additional 24 hrs. Scale bar=10 µm. Quantification of mitotic cells with multipolar spindles is provided on the right panel. (**I**) Correlation of PLK4 expression with CA20 score in breast cancer patients from METABRIC. (**J**) Expression of PLK4 in different breast cancer subtypes in METABRIC dataset. (**K**) TACC3 mRNA expression in breast cancer tumors with low vs. high PLK4 expression. (**L**) Survival analyses in breast cancer patients based on PLK4 and TACC3 expression. CC, centrosome clustering.

To experimentally demonstrate the role of TACC3 in CC, we selected two breast cancer cell lines with CA (the HER2+ cell line, JIMT-1 and the TNBC cell line, MDA-MB-231), and two non-CA cell lines (the luminal A cell line, MCF-7 and the TNBC cell line, MDA-MB-468) as negative controls (**Fig. 2B**). While TACC3 inhibition in CA cells caused an increase in the percentage of multipolar cells with scattered centrosomes (**Fig. 2C, D**), there was no increase in multipolar cell population in the non-CA cell lines, MCF-7 and MDA-MB-468 (**fig. S3**). Importantly, induction of CA with the cytokinesis inhibitor, cytochalasin D^9,22^, in MDA-MB-468 cells resulted in increased sensitivity (IC50 change from 9.4 µM to 2.4 µM) to BO-264 (**Fig. 2E**) by causing de-clustering of the extra centrosomes generated upon cytochalasin D treatment (**Fig. 2F**). Overexpression of PLK4 is another known inducer of CA which causes centrosome overduplication^23^. Overexpressing PLK4 induced CA in MDA-MB-468 cells, and TACC3 inhibition in these cells led multipolar mitosis and better cell growth inhibition (**Fig. 2G, H**), replicating the results with cytochalasin D. To demonstrate the clinical association of TACC3 with CA that is caused by PLK4 overexpression, we analyzed the METABRIC dataset^21^ and showed that PLK4, whose expression correlates with CA20 score (**Fig. 2I**), is expressed at higher levels in HER2+ and basal subtypes (**Fig. 2J**). Patients with high PLK4 expression have higher TACC3 mRNA (**Fig. 2K**) and have drastically worse overall survival (**Fig. 2L**), further validating the clinical relevance of TACC3 in cancers with CA.

### KIFC1 is a novel TACC3 binding protein that is involved in TACC3-mediated CC

Having demonstrated the phenotypic effects of TACC3 inhibition on CC, we next sought to identify the molecular mechanisms of TACC3-mediated CC in cancer cells with CA. The kinesin-14 family member, KIFC1 is among the few known CC-mediating proteins^24^. We first examined if there is a correlation between TACC3 and KIFC1 in clinical samples. We demonstrated that TACC3 expression strongly positively correlates with *KIFC1* mRNA which is also correlated well with the CC score in breast cancer patients with CA in the METABRIC dataset (**Fig. 3A, B**). Notably, in breast cancer patients with CA, combined expression of high TACC3 and high KIFC1 is more strongly associated with worse distant relapse-free survival (DRFS) compared to that of individual genes (**Fig. 3C**), suggesting that KIFC1 might work together with TACC3 in mediating CC in the CA cells. To experimentally test this hypothesis, we first examined a potential interaction between the two proteins and showed co-localization of TACC3 and KIFC1 at the centrosomes of mitotic cells by immunofluorescence staining (**Fig. 3D**). We further validated the binding between the two proteins in mitotic cells by a proximity labelling assay where we cloned TACC3 to the C-terminus of a modified peroxidase enzyme, APEX2^25^. We confirmed the spindle and centrosome localization of APEX2-TACC3 in mitotic cells (**fig. S4A**). Immunoprecipitation of TACC3 in APEX2-TACC3 overexpressing mitotic JIMT-1 cells revealed a strong binding to KIFC1 (**Fig. 3E**) in addition to its known interactor clathrin^26^. This is also verified upon H2O_2_ administration to induce specific biotinylation of the interactors, followed by streptavidin pulldown and immunoblotting for KIFC1 and ch-TOG, another known interactor of TACC3^27^ (**Fig. 3F, G, fig. S4B**). We further validated the binding between endogenous TACC3 and KIFC1 proteins by immunoprecipitating endogenous TACC3 in JIMT-1 cells (**Fig. 3H**). Importantly, the TACC3-KIFC1 interaction was abrogated upon BO-264 treatment, along with the known interactor, clathrin, as a potential molecular mechanism of TACC3 inhibition-mediated centrosome de-clustering (**Fig. 3H**). Supporting this, silencing KIFC1 with an siRNA phenocopied the effects of TACC3 knockdown in terms of the formation of multipolar spindles (**Fig. 3I-K**). To identify the KIFC1-binding region on TACC3, we transfected HEK293T cells with the full length TACC3 (1-838 aa), N-terminal (1-593 aa) and C-terminal (594-838 aa) of TACC3. These vectors express GFP and also shTACC3 to silence endogenous TACC3 protein^28^. KIFC1 IP followed by immunoblotting with a GFP antibody revealed that the C-terminal region that corresponds to the TACC domain primarily interacts with KIFC1 (**Fig. 3L, M**). Overall, these data demonstrate that (i) KIFC1 and TACC3 levels strongly correlate with each other as well as with the CC score and patient survival in the context of CA, (ii) KIFC1 is a novel interactor of TACC3 in mitotic cells, and (iii) the TACC3-KIFC1 complex mediates CC and cell survival in cancer cells with CA.

**Figure 3.**
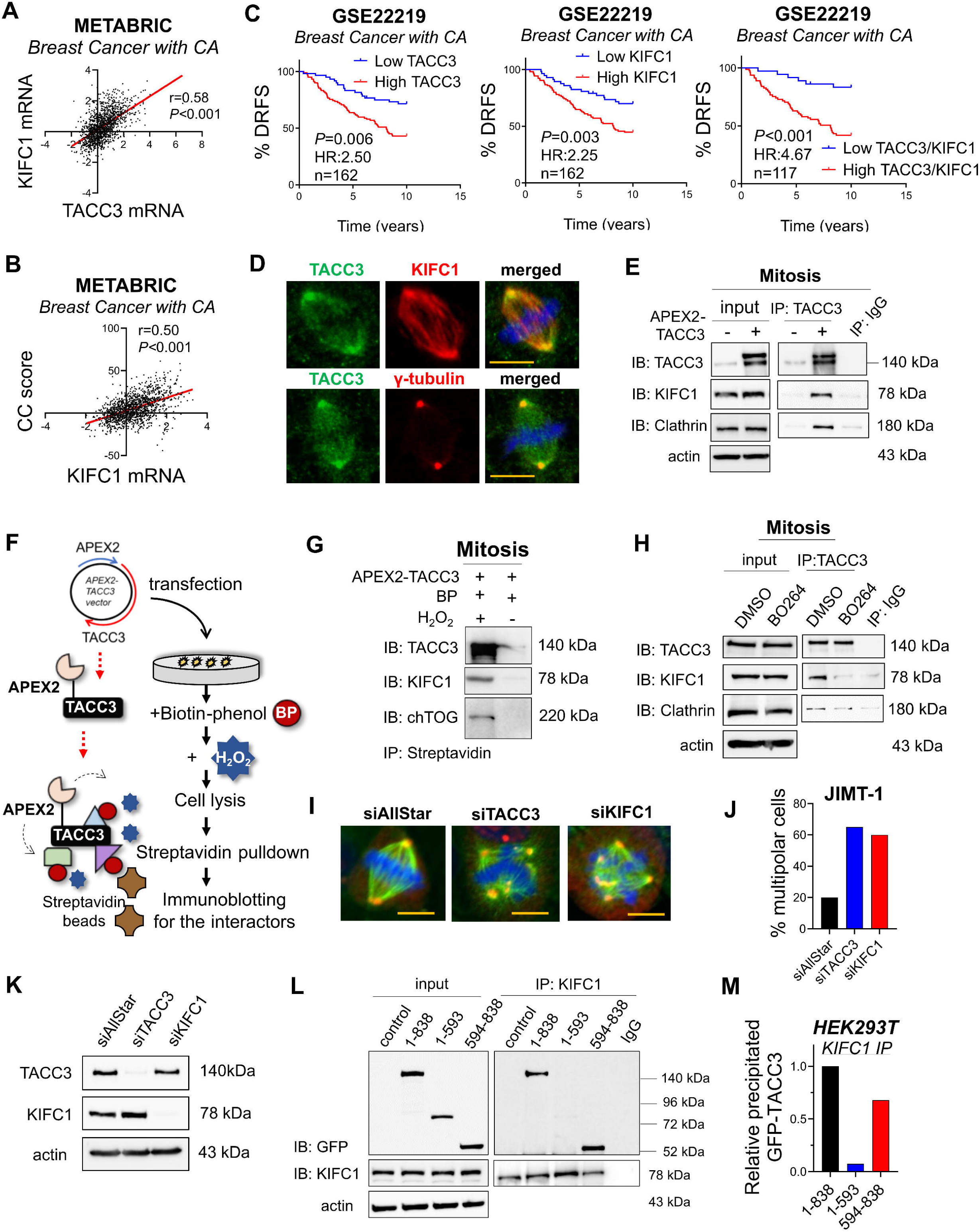
KIFC1 is a novel TACC3 binding protein that is responsible for TACC3-mediated CC. (**A, B**) Correlation of KIFC1 mRNA expression with TACC3 (A) and CC score (B) in breast cancer patients with CA in METABRIC dataset. (**C**) Survival analyses in breast cancer patients with CA based on TACC3 or KIFC1 or their combination in GSE22219 dataset. (**D**) IF staining of TACC3 (green)/KIFC1 or γ-tubulin (red) in JIMT-1 cells, showing colocalization at the centrosomes. Scale bar = 10 µm. (**E**) Western blotting of KIFC1 and the known interactor, clathrin upon TACC3 pulldown in APEX2-TACC3 overexpressing mitotic JIMT-1 cells. (**F**) APEX2 methodology as a proximity labeling technique to show binding of TACC3 to its interactors. (**G**) Western blot analysis of KIFC1 and the known interactor, chTOG upon biotinylation by H_2_O_2_ in mitotic JIMT-1 cells followed by Streptavidin pulldown. (**H**) Co-IP of endogenous TACC3 and its interactors, KIFC1 and clathrin in JIMT-1 cells synchronized at mitosis and treated with BO-264 for 4 hrs (5 µM). (**I**) IF staining of α-(green) and γ-(red) tubulin in JIMT-1 cells transfected with siTACC3 or siKIFC1 (100 nM) for 48 hrs. Scale bar = 10 µm. (**J**) Quantification of mitotic cells with multipolar spindles from I. (**K**) Western blot analysis of TACC3 and KIFC1 to validate the siRNA-mediated knockdowns. (**L**) IP of KIFC1 and IB with GFP antibody in mitotic HEK293T cells transfected with GFP-labelled vectors of different regions of TACC3. 1-838: full length, 1-593: N-terminus, 594-838; C-terminus. (**M**) Quantification of the band intensities from L. DRFS: distant relapse-free survival.

### High TACC3 expression is associated with worse survival in patients with mutant p53 and TACC3 inhibition activates the SAC/CDK1/Bcl2 axis in p53 loss/mutated cells with CA

p53 alterations are among the major causes of CA in tumors^5^. We observed that cell lines with mutant p53 (mut-p53) exhibit CA (**Fig. 1E**) and are more sensitive to the TACC3 inhibitor, BO-264 (**Fig. 4A**). To examine the clinical relevance of TACC3 in association with p53 mutations, we analyzed the METABRIC dataset by stratifying patients based on their p53 mutation status. Breast tumors with mut-p53 exhibit significantly higher levels of the CA20 score and TACC3 mRNAs (**Fig. 4B, C**), and TACC3 expression strongly correlates with CC score in mut-p53-bearing tumors (**Fig. 4D**). These results were further validated in prostate, lung and head & neck cancer patients (**fig. S5**). Notably, in breast cancer patients with mut-p53, high TACC3 expression is strongly associated with worse overall survival (**Fig. 4E**).

**Figure 4.**
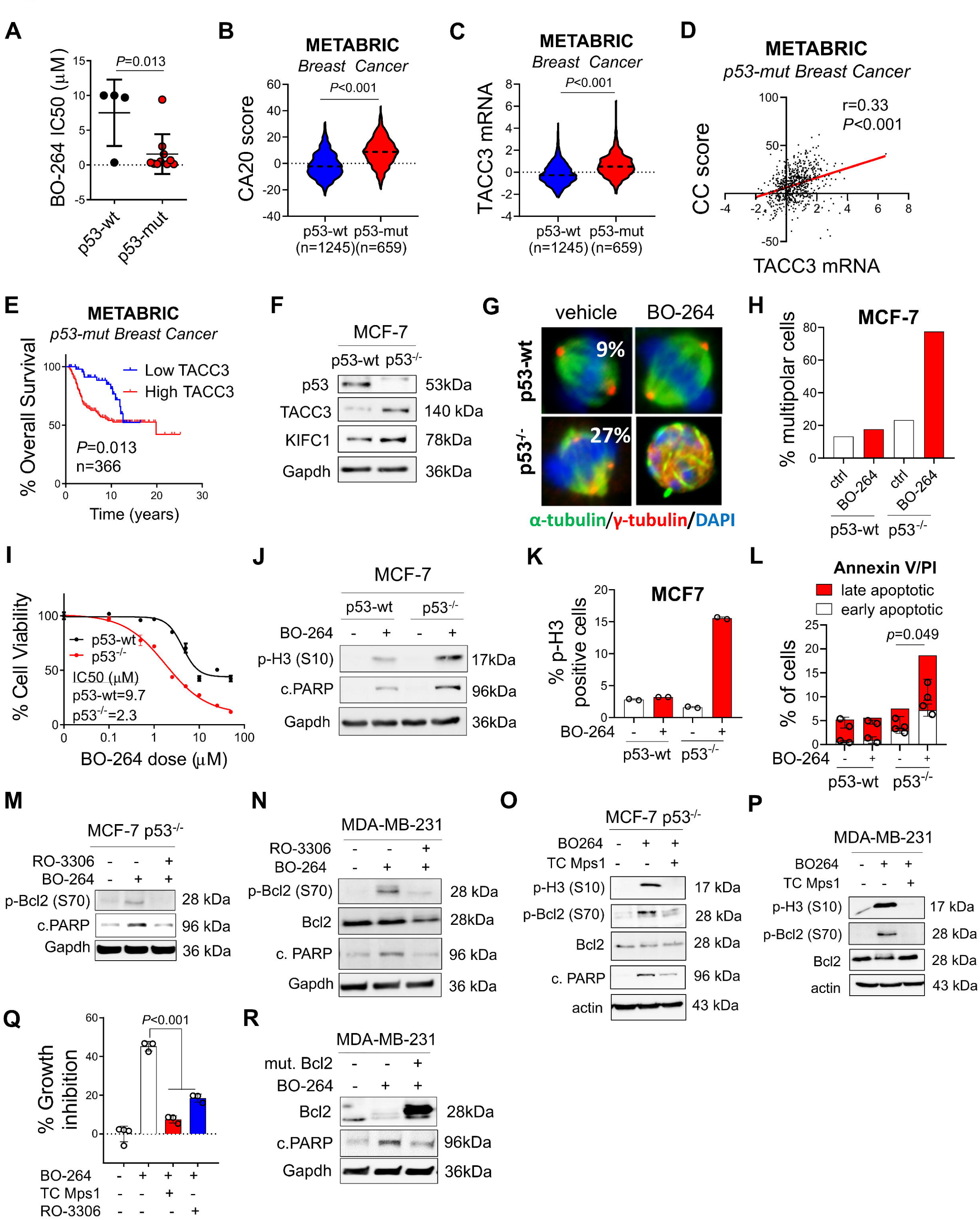
High TACC3 expression is associated with worse survival in patients with mutant p53 and TACC3 inhibition activates the SAC/CDK1/Bcl2 axis in p53 loss/mutated cells with CA. (**A**) BO-264 IC50 values in breast cancer cell lines from Fig. 1E, separated based on their p53 mutational status. (**B, C**) CA20 score (B) and TACC3 (C) expression in p53-wt vs. p53-mut breast cancer patients in METABRIC dataset. (**D, E**) Correlation of TACC3 with CC score (D) and survival analyses (E) in p53-mut breast cancer patients based on TACC3 expression in METABRIC dataset. (**F**) Western blot analysis of p53, TACC3 and KIFC1 in MCF-7 p53-wt vs p53^-/-^ cells. (**G**) IF staining of α-(green) and γ-(red) tubulin in MCF-7 p53-wt vs p53^-/-^ cells treated with 5 µM of BO-264. (**H**) Quantification of mitotic cells with multipolar spindles in cells from G. (**I**) Dose response curve of MCF-7 p53-wt vs p53^-/-^ cells to increasing doses of BO-264 for 72 hrs. (**J**) Western blot analysis of the mitotic arrest marker, p-Histone H3 (S10) and apoptosis marker, cleaved PARP in MCF-7 WT vs. p53^-/-^ cells treated with 2 µM of BO-264. (**K, L**) Flow cytometry analysis with DAPI staining (K) and Annexin V/PI staining (L) in cells from J. (**M, N**) Western blot analysis of p-Bcl2 (S70), Bcl2 and cleaved PARP in MCF-7 p53^-/-^ (M) and MDA-MB-231 (N) cells treated with BO-264 with or without 5 µM of the CDK1 inhibitor, RO-3306. (**O**) Western blot analysis of p-H3, p-Bcl2 (S70), Bcl2 and cleaved PARP in MCF-7 p53^-/-^ cells treated with BO-264 with or without 2 µM of the SAC kinase, Mps1 inhibitor, TC Mps1. (**P, Q**) Western blot analysis of p-H3, p-Bcl2 (S70) and Bcl2 (P) and percent growth inhibition (Q) in MDA-MB-231 cells treated with BO-264 with or without 2 µM TC Mps1. (**R**) Western blot analysis of Bcl2 and cleaved PARP in MDA-MB-231 cells overexpressing phospho-defective Bcl2 vector (S70A) and treated with BO-264.

To experimentally test the effects of p53 loss on CA and identify the molecular determinants of sensitivity to TACC3 inhibition in cancer cells with p53 loss or mutation, we first used p53 null (p53^-/-^) derivatives of the non-CA cell line, MCF-7 which was generated via CRISPR-Cas9 (**Fig. 4F**). Percentage of cells with CA increased to 27% from 9%, and TACC3 levels were upregulated upon p53 loss, validating the causal role of p53 in CA (**Fig. 4F, G**). Importantly, KIFC1, the novel binding partner of TACC3 that we identified, was also upregulated upon p53 loss (**Fig. 4F**). TACC3 inhibition caused centrosome de-clustering, multipolar mitosis and a stronger reduction in cell viability in p53^-/-^ cells compared to p53-wt cells (**Fig. 4G-I**). In line with this, we observed a stronger mitotic arrest and apoptosis induction in the p53^-/-^ cells compared to p53-wt counterparts upon TACC3 inhibition (**Fig. 4J-L, fig. S6**).

To decipher the molecular mechanisms of TACC3 inhibition-mediated mitotic cell death in p53^-/-^ or p53-mut cells, we examined the SAC/CDK1-dependent inhibitory phosphorylation of Bcl2^19^. Inhibiting CDK1 or the SAC kinase MPS1 in the p53^-/-^ or p53-mut cells decreased TACC3 inhibition-induced Bcl2 phosphorylation, apoptosis, and rescued cell viability (**Fig. 4M-Q**). Overexpressing a phospho-defective version of Bcl2 (S70A) in MDA-MB-231 cells reduced BO-264-induced PARP cleavage (**Fig. 4R**), further demonstrating the role of Bcl2 phosphorylation on cell death upon TACC3 inhibition in the absence of wt p53.

### TACC3 interacts with the members of the nucleosome remodeling and deacetylase (NuRD) complex in interphase cells with CA and inhibition of this interaction leads to G1 arrest and apoptosis

Although TACC3 has been shown to be a major mitotic protein, it may also have interphase functions. For instance, its inhibition has been shown to be involved in p53-dependent G1 arrest^29^; however, its mechanism and its role in p53-mut/loss cells are unknown. Interestingly, when we treated the p53 mut/CA cells JIMT-1 and MDA-MB-231 with increasing doses of BO-264, we observed a prominent increase in the CDK inhibitors, p21 (wt p53 target) and p16 and a decrease in G1/S progression markers, CDK2, cyclin D1 and RB phosphorylation (**Fig. 5A**), suggesting that in addition to its roles in mitosis, TACC3 may also play a role in G1/S progression in cancer cells with CA. Notably, when CA is induced in the non-CA MDA-MB-468 cells by treatment with cytochalasin D, cells underwent stronger G1 arrest upon TACC3 targeting (**Fig. 5B**), further suggesting a CA-driven dependence of cells to TACC3 for G1/S progression. Intriguingly, the induction of tumor suppressors upon TACC3 inhibition in CA cells was not only at protein level, but also at RNA level (**Fig. 5C, D**). These results suggest that TACC3 is involved in transcription of key tumor suppressors in interphase cells with CA in a p53-independent manner, which potentially leads to G1/S progression.

**Figure 5.**
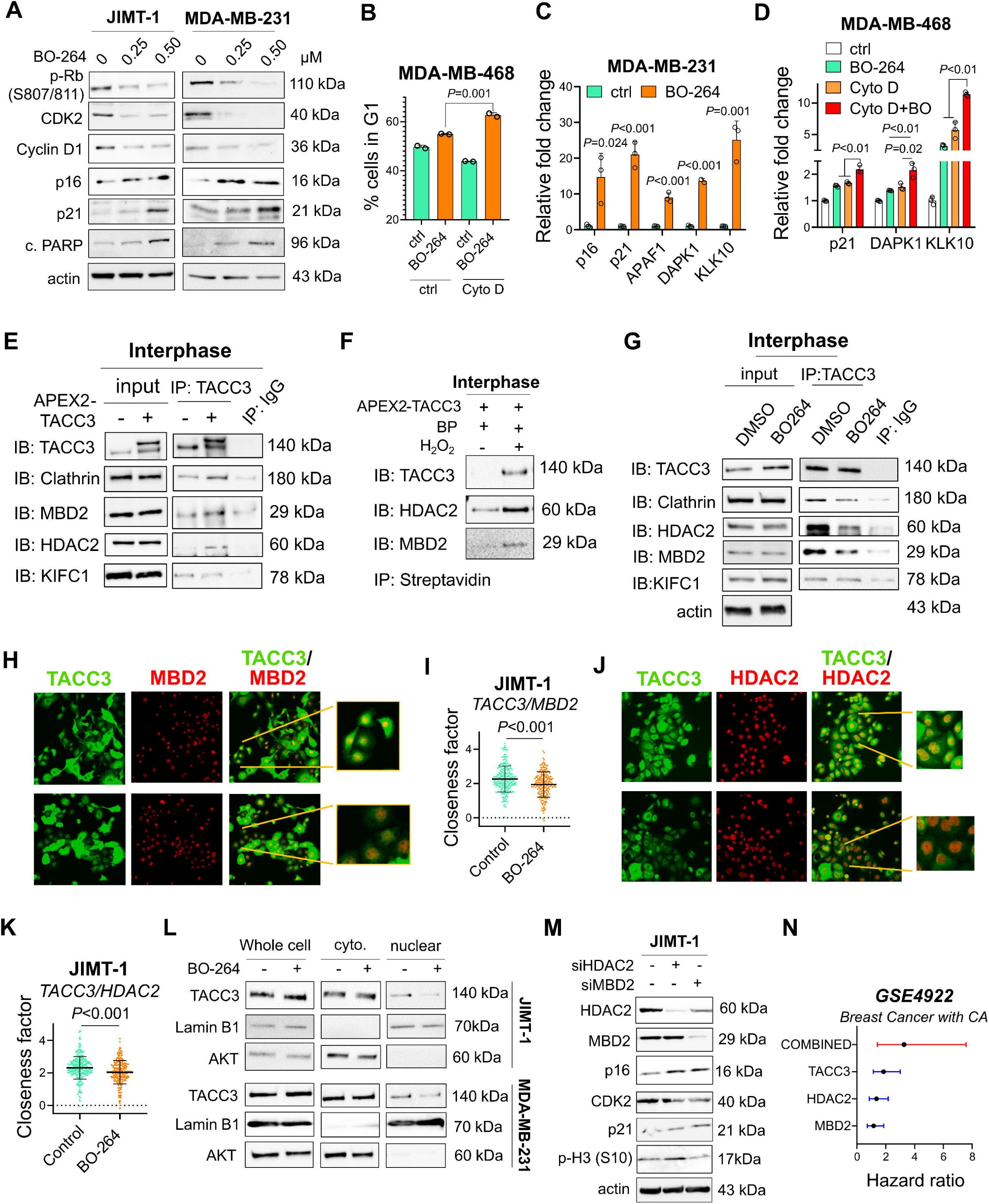
TACC3 interacts with the members of the nucleosome remodeling and deacetylase (NuRD) complex in interphase cells and inhibition of this interaction leads to G1 arrest and apoptosis. (**A**) Western blot analysis of G1/S progression markers and CDK inhibitors in JIMT-1 and MDA-MB-231 cells treated with 0.25 and 0.5 µM of BO-264. (**B**) Flow cytometry analysis with DAPI staining in MDA-MB-468 cells treated with 1 µM of cytochalasin D to induce CA followed by 24 hrs of 5 µM BO-264. (**C, D**) qPCR analysis of NuRD complex targets in MDA-MB-231 cells (C) and MDA-MB-468 cells upon CA induction by cytochalasin D (D) under BO-264 treatment. (**E**) Western blot analysis of TACC3 interactors upon TACC3 pulldown in APEX2-TACC3 overexpressing interphase synchronized JIMT-1 cells. (**F**) Western blot analysis of TACC3 interactors upon biotinylation by H_2_O_2_ followed by streptavidin pulldown in interphase synchronized JIMT-1 cells. (**G**) Co-IP of endogenous TACC3 and its interactors in JIMT-1 cells synchronized at interphase and treated with BO-264 (5 µM) for 4 hrs. (**H, I**) IF staining of TACC3 (green) and MBD2 (red) in JIMT-1 cells treated with BO-264 (H) and the closeness factor showing decrease in colocalization upon BO-264 treatment (I). (**J, K**) IF staining of TACC3 (green) and HDAC2 (red) in JIMT-1 cells treated with BO-264 (J) and the closeness factor showing decrease in colocalization upon BO-264 treatment (K). (**L**) Western blot analysis of TACC3 in cytoplasmic and nuclear fractions of interphase-synchronized JIMT-1 and MDA-MB-231 cells treated with 5 µM BO-264 for 6 hrs. Lamin B1 and AKT were used as nuclear and cytoplasmic markers, respectively. (**M**) Western blot analysis of G1/S progression markers, CDK inhibitors and mitotic arrest marker, p-H3 in JIMT-1 cells transfected with siHDAC2 and siMBD2. (**N**) Hazard ratio plot in breast cancer patients with CA based on TACC3, HDAC2 and MBD2 expressions alone or in combination using the GSE4922 dataset.

The methyl-CpG-binding protein, MBD2 is a member of NuRD complex that has important roles in processes, such as transcription, chromatin assembly, cell cycle progression and genomic stability^30^. MBD2 has been previously shown to interact with TACC3 to reactivate the transcription of methylated genes^31^. Using the APEX2 proximity labeling assay, followed by TACC3 or streptavidin IP in interphase cells, we demonstrated that MBD2 interacts with TACC3 in CA cells (**Fig. 5E, F, fig. S7A**). Nuclear localization of APEX2-TACC3 in interphase cells is validated by immunofluorescence **(fig. S7B)**. Importantly, we identified the histone deacetylase 2 enzyme (HDAC2), which is another member of the NuRD complex^32^, as a novel interactor of TACC3 in interphase cells with CA (**Fig. 5E, F**). We further demonstrated the interaction between endogenous TACC3, MBD2 and HDAC2 in interphase cells which is reduced upon treatment with TACC3 inhibitor (**Fig. 5G**). Immunofluorescence imaging showed colocalization of TACC3 with MBD2 and HDAC2 within the nucleus of CA cells which was significantly reduced upon BO-264 treatment (**Fig. 5H-K**). Interestingly, the percentage of cells with nuclear TACC3 was considerably higher in CA cell line, JIMT-1 compared to the one in non-CA cell line, MCF-7 (**fig. S8**). The nuclear localization of TACC3 was further validated by fractionation assay in interphase of two different CA cell lines, JIMT-1 and MDA-MB-231 (**Fig. 5L**). Moreover, TACC3 inhibitor, BO-264 reduced the nuclear levels of TACC3 that could explain the decrease in TACC3 interaction with MBD2 and HDAC2 upon BO-264 treatment. Furthermore, APAF1, DAPK1 and KLK10, which are among the known targets of the NuRD complex^33,34^, regulating cell cycle progression and apoptosis along with p21 and p16 were also transcriptionally activated upon BO-264 treatment in the CA cell line, MDA-MB-231 as well as in MDA-MB-468 cells upon CA induction with cytochalasin D (**Fig. 5C, D**). Notably, inhibiting HDAC2 or MBD2 with siRNAs in JIMT-1 cells with CA partially recapitulated the effects of TACC3 inhibition on the expression of G1/S transition markers without causing mitotic arrest (**Fig. 5M**). Likewise, we observed a very weak interaction of KIFC1 with TACC3 in interphase cells with no reduction upon TACC3 inhibition (**Fig. 5E, G**). These data overall suggest the presence of distinct TACC3 interactomes functioning in different cell cycle phases in cancer cells with CA. Lastly, we observed the highest hazard ratio in patients with higher expression of TACC3/MBD2/HDAC2 in combination compared to single gene expressions in breast cancer with CA in the GEO dataset, GSE4922 (**Fig. 5N**).

### Targeting TACC3 inhibits the growth of centrosome-amplified breast tumors

To test the effects of TACC3 inhibition on the growth of tumors with CA *in vivo*, we performed CRISPR Cas9-mediated knockout of TACC3 (named sgTACC3) in JIMT-1 and MDA-MB-231 cells and validated the TACC3 depletion by Western blotting (**Fig. 6A, C**). TACC3 knockout significantly decreased the colony formation ability of the cells (**Fig. 6B, D**). To test whether the TACC domain of TACC3, which we found to be important for KIFC1 binding and also known to be required for MBD2 interaction^31^, is able to increase the transforming ability of normal cells, we transfected the normal breast cells, MCF12A with full length, N- and C-terminal domains of TACC3. As shown in **Fig. 6E**, the TACC domain (594-838) was sufficient to induce colony formation to a similar extent to full length, whereas the N-terminal region (1-593) failed to increase the transforming ability.

**Figure 6.**
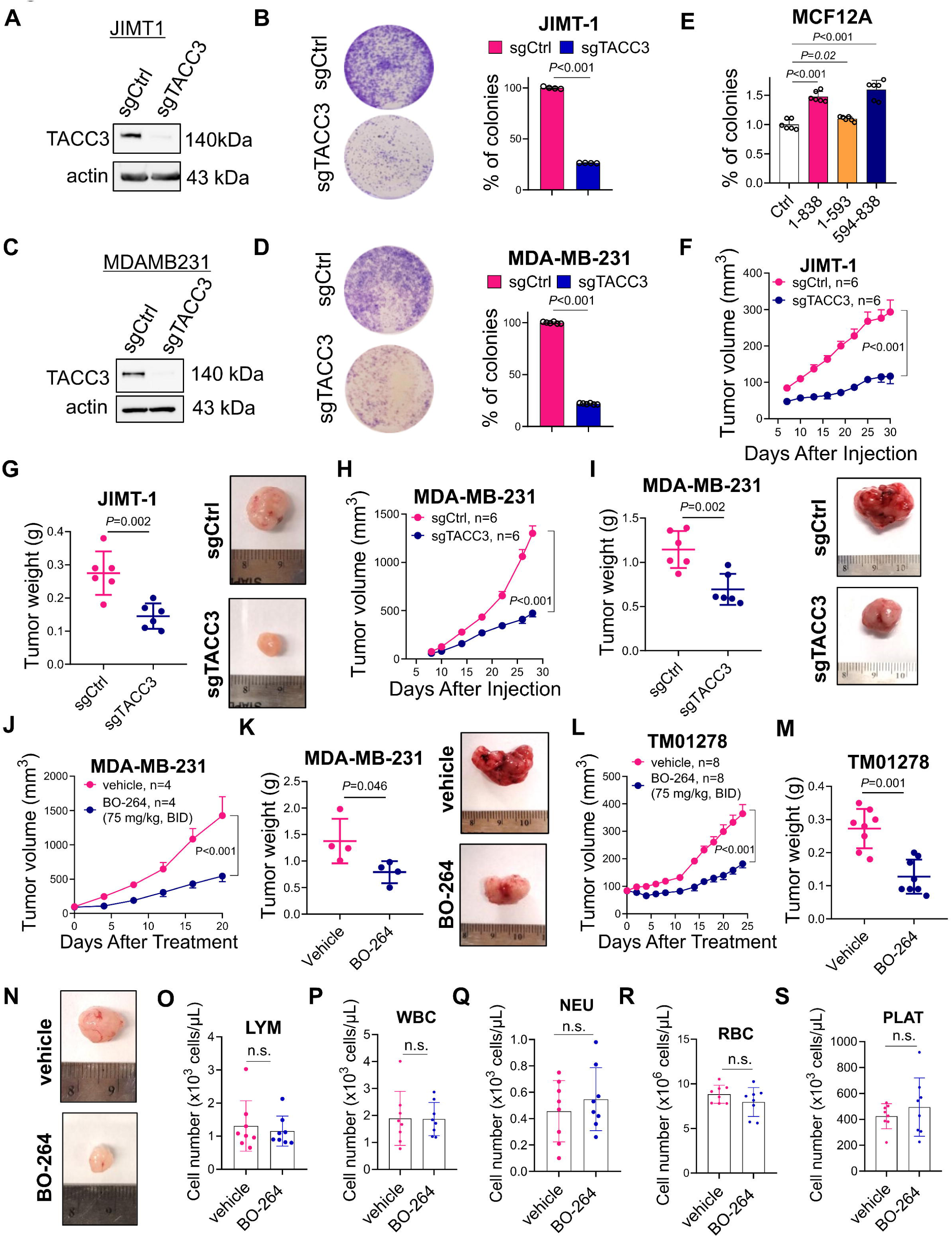
Targeting TACC3 inhibits tumor growth in centrosome-amplified breast tumors *in vivo*. (**A-D**) Western blot validation of CRISPR/Cas9-mediated knockout of TACC3 (A, C) and the effect of TACC3 knockout on colony formation (B, D) in JIMT-1 and MDA-MB-231 cells. (**E**) Relative colony formation ability of MCF12A cells overexpressing different regions of TACC3. 1-838: full length, 1-593: N-terminus, 594-838; C-terminus. (**F, G**) Tumor growth (F) in xenografts of JIMT-1 sgCtrl vs. sgTACC3 cells, and the tumor weights and representative images at the end of the experiment (G). (**H, I**) Tumor growth (H) in xenografts of MDA-MB-231 sgCtrl vs. sgTACC3 cells, and the tumor weights and representative images at the end of the experiment (I). (**J**) Tumor growth in xenografts of MDA-MB-231 cells treated with 75 mg/kg BO-264, twice daily (p.o.). (**K**) The tumor weights and representative images from vehicle vs. BO-264 treated mice from J at the end of the experiment. (**L**) Tumor growth of TNBC PDXs, TM01278 treated with 75 mg/kg BO-264, twice daily (p.o.). (**M, N**) The tumor weights (M) and representative images (N) from vehicle vs. BO-264 treated mice from L in the end of the experiment. (**O-S**) Blood cell counts of the mice from L at the end of the experiment.

To test the effects of TACC3 knockout on *in vivo* tumor growth, we injected sgCtrl vs. sgTACC3-expressing JIMT-1 and MDA-MB-231 cells to the mammary fat pad (MFP) of nude mice and monitored tumor growth. As shown in **Fig. 6F-I**, TACC3 knock-out reduced the growth of tumors with CA. In addition, we tested the effects of pharmacological inhibition of TACC3 on tumor growth in both MDA-MB-231 xenografts and the TNBC PDX, TM01278 with CA using the TACC3 inhibitor, BO-264. TACC3 inhibition significantly reduced the growth the tumors without any observable toxicity as shown by the blood cell counts and the body weights of mice (**Fig. 6J-S, fig. S9**). Altogether, these data suggest that TACC3 inhibition represents a novel vulnerability in tumors with CA.

## DISCUSSION

CA is one of the hallmarks of cancer^1,2^ and is associated with tumor aggressiveness and worse clinical outcome in many cancers, including breast, lung and prostate cancers^2^. Given the key roles of centrosomes in supporting bipolar spindle formation, CC is among the major coping mechanisms utilized by cancer cells with CA to prevent multipolar mitosis and cell death. In clinical settings, strategies targeting mitotic cells were shown to be inadequate due to small fraction of mitotic cells within the tumors^11^. Therefore, novel strategies targeting not only the CC machinery and mitosis, but also the non-mitotic cells with CA may have huge translational potential to eradicate tumors with CA without impacting the normal cells. Here, we identified the centrosome-associated protein, TACC3 as an essential gene for the survival of cancer cells with CA. We showed that TACC3 expression is higher in high CA tumors, and it significantly correlates with CC signature and worse clinical outcome in cancer patients with CA. It is also strongly correlated with p53 mutations and PLK4 expression, which are among the major causes of CA in highly aggressive tumors. Mechanistically, we showed, for the first time, that TACC3 interacts with the CC protein, KIFC1 and ensures faithful mitosis by promoting clustering of extra centrosomes in cancer cells with CA. We also uncovered novel functions of TACC3 beyond mitosis that involves interaction with members of the transcriptional regulator, NuRD complex in nucleus, and regulation of the transcription of genes responsible for G1/S transition and cell survival/apoptosis (**Fig. 7A**). Inhibiting TACC3, on one hand, blocks its interaction with KIFC1, leading to centrosome de-clustering and mitotic catastrophe via the SAC/CDK1/p-Bcl2 axis, and on the other hand induces p53-independent G1 arrest by relieving the inhibitory effect of the NuRD complex on the transcription of key tumor suppressors (**Fig. 7B**), culminating in apoptosis and inhibition of CA-driven tumor growth. Overall, our findings provide valuable preclinical data for targeting TACC3 in highly aggressive tumors bearing amplified centrosomes.

**Figure 7.**
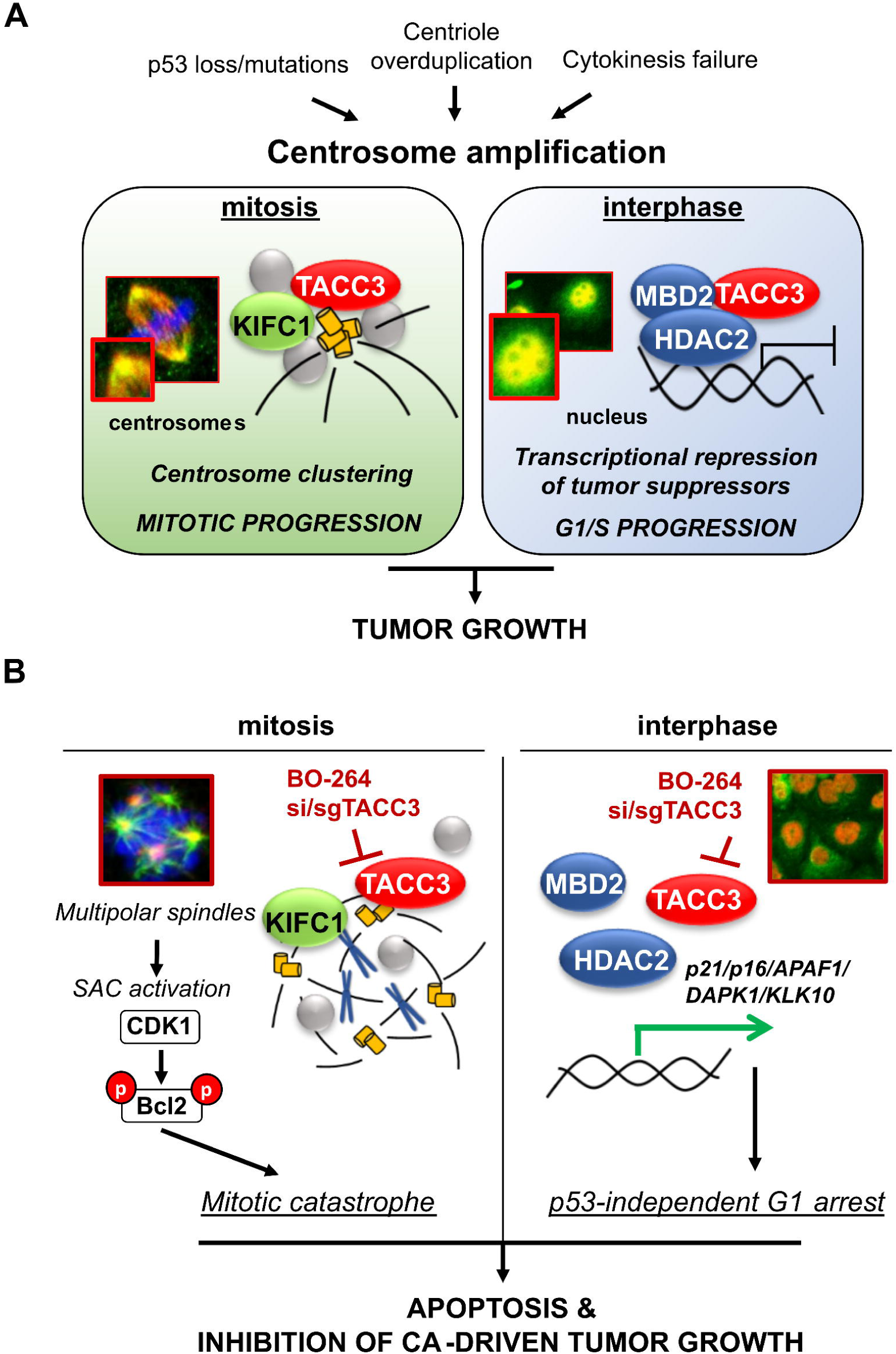
Schematic summary of the findings. (**A**) In cancers with amplified centrosomes induced via various causes, such as PLK4 overexpression, p53 mutations or cytokinesis failure, TACC3 is overexpressed and mediates distinct mitosis and interphase-specific functions. Mitotic TACC3 interacts with KIFC1 at the centrosomes and promotes CC while interphase TACC3 interacts with MBD2 and HDAC2, belonging the NuRD complex, to suppress the transcription of tumor suppressors. (**B**) Upon TACC3 targeting by BO-264 or si/sgRNAs, cells, on one hand, undergo mitotic catastrophe due to loss of TACC3-KIFC1 interaction and centrosome de-clustering, and on the other hand undergo p53-independent G1 arrest via activation of tumor suppressor transcription, culminating in apoptosis and inhibition of tumor growth.

TACC3 is known to be overexpressed in cancer, localizes to spindles and centrosomes, and required for proper cell division^17,18^. Despite being known to mediate centrosome integrity, its potential functions in cancer cells with CA and its clinical relevance in the context of CA have not been studied. Our findings demonstrate, for the first time, that TACC3 is overexpressed in cancers with CA and targeting TACC3 represents a novel vulnerability in cancer cells with CA. Aurora A, a multifaceted kinase that can phosphorylate TACC3^35^, is known to cause CA partly via phosphorylating p53 at S315, marking it for degradation^36^. Loss of p53 can, in turn, trigger CA via controling centrosome duplication^37^. Based on our novel findings that showed strong upregulation of TACC3 in p53^-/-^ or p53-mut cells/tumors, it would be interesting to test whether Aurora A induces CA by regulating TACC3. We further showed that TACC3 triggers CC in mitotic CA cells, thereby ensures bipolar spindle formation and proper mitosis. We identified a novel TACC3 interactor, KIFC1 which is known to induce spindle pole focusing necessary for the survival of cancer cells with CA^38^. We demonstrated that TACC3 binds to KIFC1 via its TACC domain at the centrosomes of mitotic CA cells and confirmed the robustness of this interaction using multiple approaches, including state-of-the-art APEX2 proximity labeling assay. Interestingly, KIFC1-mediated CC has recently been shown to involve phosphorylation by ATM/ATR kinases under DNA damaging agents, leading to drug resistance^39^. In this line, it is yet to be determined whether KIFC1 phosphorylation may be critical for the formation of the TACC3/KIFC1 complex as well, in mitotic CA cells. Future studies may also investigate if TACC3 phosphorylation by Aurora A which has been implicated in CC^40^ is required for TACC3/KIFC1-mediated CC. Furthermore, unbiased proteomic approaches can identify the unknown members of this large multiprotein complex on centrosomes which will help identify the interactome of TACC3-driven CC and may also unleash new drug targets.

TACC3 was shown to co-localize to nuclear envelope and maintain proper nuclear envelope structure^41^. It can also interact with MBD2^31^, a methyl-CpG-binding protein and a member of the NuRD complex, which regulate processes, such as transcription and chromatin assembly^30^. Although TACC3 depletion has been shown to induce G1 arrest in a p53-dependent manner^29^, the underlying molecular mechanisms in cancer cells with p53 loss or mutations that may also involve MBD2 or other NuRD members have largely been unknown. Here, we identified HDAC2, a histone deacetylating enzyme that was shown to be a part of the MBD2-containing NuRD complex^32^ as a novel interactor of TACC3 in interphase cells with CA. We showed, for the first time, that TACC3 inhibition reduces nuclear TACC3 levels, disrupts the interaction of TACC3 with the NuRD complex (i.e., MBD2 and HDAC2), and thus, relieving the inhibitory effect on the transcription of key tumor suppressors, i.e., p16, p21, APAF1, DAPK1 and KLK10 in a p53-independent manner, ultimately leading to G1 arrest and apoptosis. Among those, p21 and APAF1 are known to be direct targets of p53, suggesting that blocking NuRD complex via TACC3 inhibition can activate the transcription of tumor suppressors even in the absence of functional p53, e.g., in tumors with CA. Importantly, despite the key roles of the NuRD complex in tumor progression, there are no inhibitors specifically targeting NuRD complex. Based on our novel findings, TACC3 inhibition may represent a novel way of inhibiting NuRD complex in cancer.

A few mitotic proteins have been previously identified as potential targets for eliminating centrosome-amplified, chromosomally instable (CIN) cancer cells^44^. For instance, the mitotic kinesin, KIF18A has been shown to play a key role in maintaining bipolar spindle integrity and to be crucial for the survival of cancer cells with CIN^42^. Inhibitors against the CC-mediator, KIFC1 (e.g. AZ82^43^) have been shown to be effective *in vitro*; however, their preclinical testing in cancer models is lacking. Inhibitors against Aurora kinases have already been clinically tested in solid tumors and hematologic malignancies; however, only poor or modest efficacy was observed, along with side-effects, such as neutropenia^13^. Alisertib (MLN8237) is the only Aurora A inhibitor progressed to Phase III evaluation in Peripheral T-Cell Lymphoma; however, no superior benefit was achieved over its investigator-selected single-agent comparator^44^. Therefore, it is of utmost importance to identify more effective therapeutic strategies against cancers with CA which preferentially target not only mitotic but non-mitotic cancer cells to achieve superior clinical efficacy^11^. Along these lines, TACC3 may represent an excellent drug target with a strong translational potential in the treatment of highly aggressive cancers as 1.) it is overexpressed in CA tumors compared to non-CA tumors and normal tissue; 2.) its inhibition blocks cell cycle progression at both mitosis and interphase with distinct mechanisms, assuring effective inhibition of cell proliferation; and 3.) its inhibition has strong anti-tumor activity with no apparent toxicity *in vivo*.

We demonstrated the CA-directed vulnerability to targeting TACC3 by inducing CA in non-CA cells using different approaches, such as PLK4 overexpression or cytokinesis inhibition by cytochalasin D which conferred high sensitivity to TACC3 inhibition. Percentage of tumors with CA is profoundly high in many different cancer types, such as invasive breast carcinoma and squamous cell carcinomas of the head and neck where percentage of tumors with hyper-amplified centrosomes goes up to 80%^45^. CA is also associated with several hallmarks of cancer related to tumor aggressiveness, such as aneuploidy that is observed in the vast majority (∼70%) of solid tumors^46^, and p53 mutations that occurs in ∼50-60% of tumors, and can be seen in as high as 80% of patients of aggressive cancer subtypes, such as TNBCs^47^. Therefore, strategies targeting the CA-driven vulnerabilities has the potential to be highly effective in aggressive tumor types where the percentage of cells with CA or the associated phenotypes is extremely high. Importantly, p53 mutations as well as amplified centrosomes were also shown to positively associate with tumor aggressiveness and development of metastatic disease^48^. Considering the roles of centrosomes in directing cell polarity and movement, it is highly likely that TACC3 may also be involved in tumor cell dissemination in the context of CA and p53 mutant cancers. Along these lines, targeting cancer cells with mut-p53 and/or CA by TACC3 inhibition would be a highly effective strategy against metastatic dissemination and would further improve clinical outcome at the later stages of the disease.

Overall, we identified TACC3 as a novel CC-mediator and a transcription-regulator overexpressed in highly aggressive tumors with CA and associated with drastically worse clinical outcome. We showed, for the first time, that TACC3 forms distinct functional interactions in different phases of the cell cycle that are essential for the survival of cancer cells with CA. We demonstrated that mitotic TACC3 inhibition causes centrosome de-clustering by disrupting the novel TACC3-KIFC1 interaction at the centrosomes, leading to SAC/CDK1/p-Bcl2-dependent mitotic cell death in cancer cells with CA. In non-mitotic cells, TACC3 inhibition causes G1 arrest by inhibiting the NuRD complex in the nucleus, leading to transcription of key cell cycle/apoptosis-regulating tumor suppressors, e.g., p21, p16 and APAF1 (**Fig. 7B**). These preclinical findings as well as the supporting clinical data strongly encourage the clinical testing of TACC3 inhibitors to improve the outcome of highly aggressive cancer patients bearing amplified centrosomes.

## MATERIALS AND METHODS

### Cell culture and reagents

Human breast cancer cell lines, ZR-75-1, MDA-MB-231, MDA-MB-157, MDA-MB-436, MDA-MB-468, CAL51, HCC1954, JIMT-1, MCF-7, T47D, SK-BR-3, BT-474 and HCC1143 and the normal breast cells, MCF12A were obtained from ATCC (Manassas, VA, USA). All the cells were cultured in Dulbecco Modified Eagle Medium (Lonza, NJ, USA) supplemented with 50 U/ml penicillin/streptomycin, 1% non-essential amino acids and 10% fetal bovine serum (Lonza, NJ, USA). The media for ER+ cell lines were further supplemented with insulin (0.1 µg/ml). The cell lines were authenticated and tested for mycoplasma contamination regularly using MycoAlert mycoplasma detection kit (Lonza, NJ, USA). The cumulative culture length of cells between thawing and use in this study was less than 20 passages.

### Inhibitor treatments and cell viability assays

BO-264, cytochalasin D, TC Mps1, and RO-3306 (Sigma Aldrich, MO, USA) were dissolved in DMSO. For cell viability assays, cells were treated with the inhibitors for 48-72 hrs, and cell viability was measured by Sulforhodamine B (SRB) (Sigma Aldrich, MO, USA). Induction of CA was performed by treating cells with 1 uM of cytochalasin D for 20 hrs, followed by 24 hrs incubation in drug-free media. TC Mps1 and RO-3306 were given 6 hrs prior to collection of cell pellet at doses of 2 µM and 1 µM, respectively^49^. The mitosis and interphase synchronizations were done using 100 ng/mL nocodazole for and double thymidine block (2 mM), respectively.

### PDX-derived organoids

TNBC PDX organoids were established as previously described^50^. For drug testing studies, organoids were dissociated with Tryple (Gibco, MA, USA) at 37 ºC for 30 mins in the presence of 10µM Rock inhibitor (Selleckchem, TX, USA). After counting the single cells, they were plated into 96-well plate (20,000 cells/well) on a matrigel-coated surface with media containing 2% matrigel (Corning, NY, USA). BO-264 was added 72 hrs after seeding. Organoids were grown in the presence of drug or vehicle for 7 days, and the organoid viability was measured using 3D Cell Titer Glo (Promega, WI, USA).

### Transient transfection with siRNAs and overexpression vectors

siRNA transfections were done in P/S-free growth medium with reduced serum at a concentration of 40 nM using Lipofectamine 2000™ (Invitrogen, CA, USA) as previously described^51^. The list of siRNAs used (Dharmacon, CO, USA) are provided in **table S1**. The PLK4 and Bcl2 S70A vectors were given 24 hrs before treatment with BO-264. HEK293T cells were transfected with TACC3 truncated vectors for 24 hrs and next day synchronized at mitosis using 100 ng/mL nocodazole.

### CRISPR/Cas9-mediated knockout studies

The sgRNA sequence targeting TACC3 in JIMT-1 cells is: 5’-CAGGCAACGTACCCTCAGCG-3’, and the sgRNA sequence targeting TACC3 in MDA-MB-231 cells is: 5’-GACTTGGTGTCACCTCCGAA-3’. sgRNAs were designed and selected based on having high on-target (=high efficacy) and low off-target (=high specificity) activity using the CRISPick tool (Broad Institute). The designed sgRNAs were cloned into human lentiCRISPR v2 vector (Addgene, MA, USA). For lentiviral packaging, HEK293T cells were transfected with sgRNAs and the packaging plasmids, pMD2.G and psPAX2 (Addgene, MA, USA). Transduction of JIMT-1 and MDA-MB-231 cells was performed in the presence of 10 ug/ml polybrene, and selection of transduced cells was done using 2µg/ml puromycin.

### APEX2 proximity labeling

JIMT-1 cells were transfected with APEX2-TACC3 vector and synchronized for mitosis and interphase as described above. Biotinylation was performed as previously described^52^. Briefly, cells were treated with 2.5 mM biotin phenol (Iris Biotech, Germany) for 1 hr, and 1 mM H_2_O_2_ was added at room temperature (RT) for 2 min. After washing, cells were lysed with RIPA buffer + quenchers (5mM Trolox, 10 mM NaN_3_, and 10mM Sodium Ascorbate). Lysates were sonicated and clarified by centrifugation. Pre-washed Dynabead M-280 Streptavidin beads were incubated with the cell lysate at 4°C for 4 hrs. After the incubation, beads were washed and boiled in elution buffer, and samples were loaded onto polyacrylamide gel for immunoblotting the interactors.

### Colony formation assay

Single-cell suspensions of sgCtrl and sgTACC3-expressing JIMT-1 and MDA-MB-231 cells (3×10^3^ cells/well) were plated in a 12-well plate. After 1 week of seeding, cells were fixed with 4% paraformaldehyde for 30 min and stained with 1% crystal violet (Merck, Darmstadt, Germany) for 20 min at RT. The plates were air dried, and the dye was dissolved in 10% methanol and 10% acetic acid mixture followed by measuring absorbance at 590 nM. Quantification of percentage colonies was done by subtracting background reading of the reagent from the all the readings, followed by normalization to sgCtrl expressing cells.

### Quantitative RT-PCR analysis

Total RNA isolation, cDNA synthesis and quantitative real-time PCR assay were performed as previously described^19,50^. The sequences of the qRT-PCR primers are provided in **table S2**. For data analysis, ΔΔ*C*_T_ method was utilized using Excel (Microsoft, WA, USA).

### Western blotting

Protein isolation and Western blotting were done as previously described^49^. Briefly, proteins were extracted using RIPA lysis buffer with the addition of protease and phosphatase inhibitor cocktails. Equal amounts of protein were separated using 10% SDS-PAGE gel. Separated proteins were transferred to PVDF membranes (Bio-Rad, CA, USA) using a Trans-Blot turbo transfer system (Bio-Rad, CA, USA) and incubated with primary antibodies (**table S3**) overnight at 4°C followed by secondary antibody incubation, and signal detection by chemiluminescence. Images were acquired using Image Lab Software (Biorad, CA, USA).

### Immunoprecipitation

After synchronizing JIMT-1 cells in mitosis or interphase, cells were treated with 5 µM of BO-264 for 4 hrs. Cells were lyzed in lysis buffer (50 mM TrisHCl pH=7.0, 150 mM NaCl, 0.2% NP40, 7.5% glycerol, protease and phosphatase inhibitor cocktail), and clarified by centrifugation. 1 mg protein for each condition was incubated with antibody-coated Dynabeads Protein G (Invitrogen, 1003D). Antibody dilutions are provided in **table S3**. Beads were washed and resuspended in 60 µL 1X SDS sample loading buffer, boiled for 10 min at 70 °C and loaded onto polyacrylamide gel.

### Immunofluorescence and quantification

Immunofluorescence staining was performed as previously described^19,50^. Dilutions for the primary and secondary antibody incubations are provided in **table S3**. Images were taken with Zeiss LSM700 Confocal Microscopy and analyzed using ImageJ software^53^. Quantification of cells with multipolar or bipolar spindles was done by counting the number of mitotic cells with >2 scattered centrosomes (multipolar) or with 2 or more centrosomes clustered in a bipolar manner (bipolar) as determined by staining centrin 2 or γ-tubulin as centrosomal markers. Then, the ratio to total number of mitotic cells was calculated and represented as percentage. At least 100 cells for treatment groups and 20-50 cells for the control groups were quantified. Quantification of the colocalization between TACC3 and MBD2/HDAC2 was done by calculating the closeness factor as defined by the log transformed value of 1/[incremental change in the ratio of the intensities of TACC3 and HDAC2/MBD2]. At least 200 cells were quantified for the colocalization analysis.

### Annexin V/PI staining and cell cycle assay

Annexin V/PI staining was done as previously described^19,50^. Briefly, counted cells for each condition were washed with PBS and incubated with 1.5 µL of FITC-conjugated Annexin V and PI for 30 min. Data collection was done with BD FACSDiva software (BD, NJ, USA), and analysis with De Novo FCS Express software. For cell cycle analysis, cells were fixed and permeabilized in 70% ethanol and then incubated with p-Histone H3 antibody and anti-rabbit Alexa-Fluor 488 antibody for 45 min at RT, sequentially. Lastly, cells were incubated in DAPI for 15 min at RT followed by data collection and analysis.

### *In vivo* studies

Six-to-eight-week-old female BALB/c nude or Nu/J mice were housed with a temperature-controlled and 12-hour light/12-hour dark cycle environment. All the *in vivo* studies were carried out in accordance with the Institutional Animal Care and Use Committee of the University of South Carolina. The sgCtrl and sgTACC3-expressing derivatives of MDA-MB-231 and JIMT-1 cells were injected into MFPs of female BALB/c nude or Nu/J mice at a cell number of 5×10^6^ and 4×10^6^ cells, respectively, in 100 μl of 1:1 PBS and Matrigel (Corning, NY, USA), v/v). Primary tumor growth was monitored by measuring the tumor volume at least twice a week with a digital caliper. Tumor volumes were calculated as length × width^2^/2. All mice used were of the same age and similar body weight.

For testing the effects of TACC3 inhibitor, BO-264 on tumor growth, 5×10^6^ MDA-MB-231 cells were injected into MFPs of female BALB/c nude mice. Once the mean and median of tumor volume reached around 100 mm^3^, xenografts were randomized into two groups (4 mice per group). Animals were treated with vehicle or BO-264 (twice a day with 75 mg/kg oral gavage (po.)). The PDX experiment was carried out by transplanting 2×2×2 mm to 3×3×3 mm sized fragments to the flank region of female Nu/J mice. Once the mean and median of tumor volume had reached around 85 mm^3^, PDXs were randomized into two groups (8 mice per group), and treated with vehicle, or BO-264 (twice a day with 75 mg/kg, po.).

### Bioinformatics and statistical analyses

The METABRIC data^21^ were downloaded from EMBL European Genome–Phenome Archive (http://www.ebi.ac.uk/ega/) with an accession number EGAS00000000122. The Cancer Genome Atlas (TCGA)^54^ patient data (prostate cancer) and the CCLE data^55^ were downloaded using the cBIO dataset^56^. The microarray data sets, GSE25066^57^, GSE4922^58^, GSE22219^59^, GSE31210^60^ and GSE41613^61^ were download from the GEO database. The CA20^20^ and CC^9^ signature scores were calculated by summing up the z-scores of the genes found in the list^62^ using the SPSS Statistics software. Seventy five percent of breast cancer patients^63^ and 83% of head & neck cancer patients^45^, 50% of lung^64^ and prostate^65^ cancer patients were stratified as high CA based on the CA20 score expression. Cancer patients except breast cancer were separated as low vs high TACC3 expressers based on median expression, and breast cancer patients were separated as low vs high TACC3 based on median or 25^th^ percentile.

The results are represented as mean□±□standard deviation (SD) or mean□±□standard error of the mean (SEM), as indicated in the figure legends. All statistical analyses were performed in GraphPad Prism Software. Comparisons between two groups were done using paired two-sided Student’s t-test for tumor growth graphs, and unpaired two-sided Student’s t-test for all other comparisons. z-score calculations were done using the SPSS Statistics software. Survival curves were generated based on median or 25th percentile separation using Kaplan-Meier method, and significance between groups was calculated by Log-rank test. Hazard ratio plot depicts the log rank hazard ratio with lower and upper 95% confidence interval. For correlation analysis, Pearson correlation coefficients were calculated.

## Acknowledgements

We are thankful to the members of Ozgur Sahin laboratory for invaluable discussion and advice. We also thank the Center for Targeted Therapeutics (CTT) Microscopy and Flow Cytometry Core (Dr. Chang-uk Lim) of the University of South Carolina, for assistance with flow cytometry. We thank METABRIC Consortium^21^ for providing us with the breast cancer gene expression profiling data. We thank Drs. Erden Banoglu and Burcu Caliskan (Gazi University) for providing us with BO-264; Dr. Stephen Royle (University of Warwick) for providing us TACC3 vectors; Dr. Phillip Buckhaults (University of South Carolina) for providing us the p53^-/-^ MCF7 cells; and Dr. Onur Cizmecioglu (Bilkent University) for providing us the APEX2 vector

## Funding

This work was supported by NIH Research Project Grant R01-CA251374 (OS) and Susan G. Komen Interdisciplinary Graduate Training to Eliminate Cancer Disparities (IGniTE-CD) GTDR17500160 (OzgeS).

## Author contributions

Ozge S. designed and performed experiments, acquired and analyzed data, interpreted data, and prepared the paper; O.A. contributed to drug response assays with TACC3 inhibitor and analyzing the NCI60 panel drug screen data; M.C. contributed to CRISPR-Cas9-mediated knockout experiments and APEX2 proximity assay; V.S. contributed to analyzing the immunofluorescence images. O.S. designed the study, oversaw experiments and data analyses, and prepared the paper. All authors reviewed and commented on the paper.

## Competing Interests

O.S. is the co-founder and manager of OncoCube Therapeutics LLC and founder and president of LoxiGen, Inc. The other authors declare no potential conflict of interest.

## Data and materials availability

All data associated with this study are present in the paper or the Supplementary Materials. Raw data of the main and supplementary figures are provided in data file S1. Gene expression data were downloaded from the NCBI Gene Expression Omnibus database under GSE31210, GSE4922, GSE22219, GSE25066, and GSE41613. METBARIC data were downloaded from EMBL European Genome–Phenome Archive (http://www.ebi.ac.uk/ega/) with an accession number EGAS00000000122. The Cancer Genome Atlas (TCGA) patient data (prostate cancer) and the CCLE data were downloaded using the cBIO dataset.

## Notes

### Competing Interest Statement

Ozgur Sahin is the co-founder and manager of OncoCube Therapeutics LLC and founder and president of LoxiGen, Inc. The other authors declare no potential conflict of interest.

