## Supplementary Data for "Targeting TACC3 represents a novel vulnerability in highly aggressive breast cancers with centrosome amplification"

Ozgur Sahin, PhD

Associate Professor

University of South Carolina

Department of Drug Discovery and Biomedical Sciences

715 Sumter Street, CLS609D, Columbia, SC, 29208

**Running Title:** Targeting TACC3 in centrosome amplified breast cancer

**Keywords:** Centrosome amplification/centrosome clustering/TACC3/KIFC1/NuRD complex

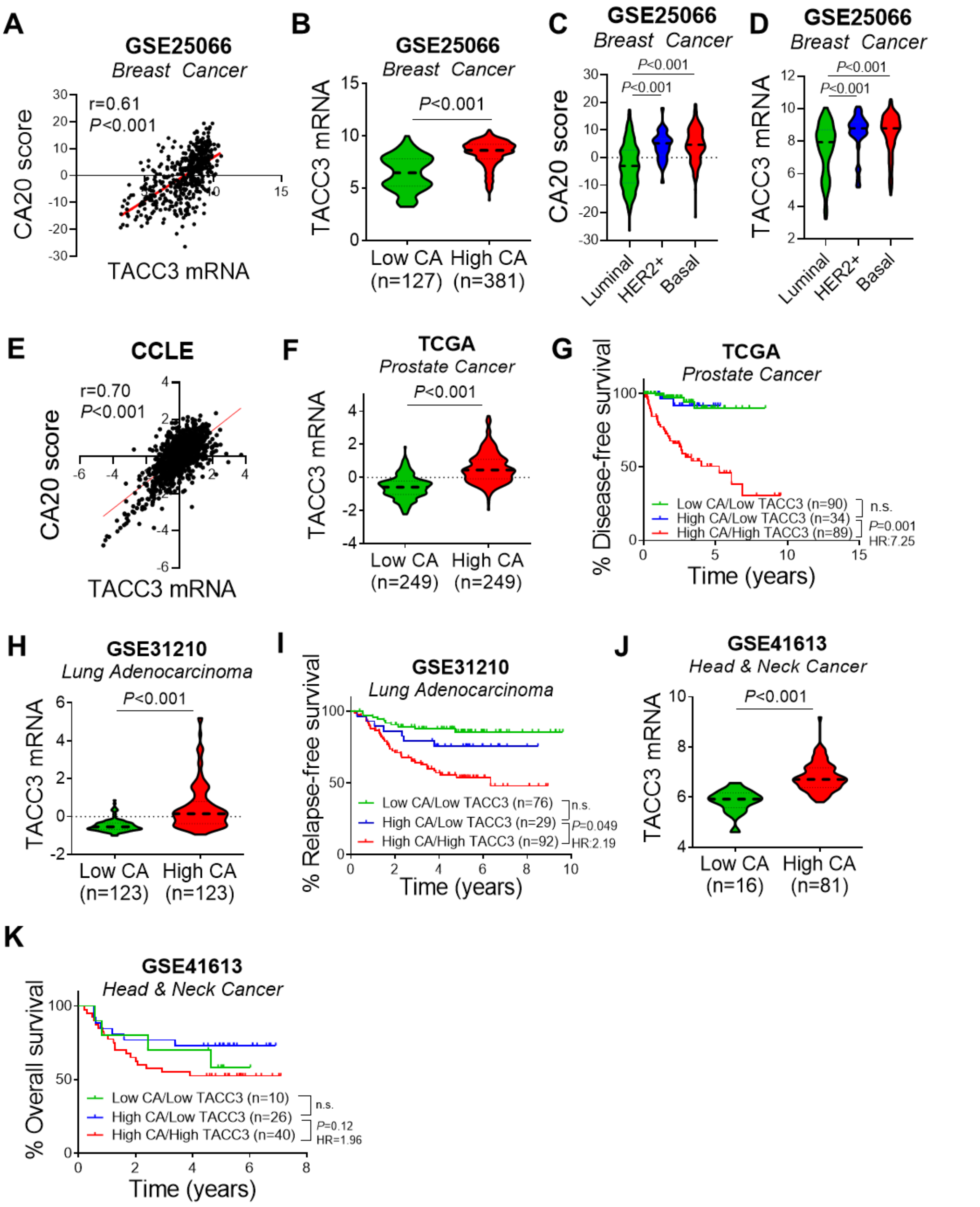


**Fig. S1. TACC3 is upregulated in highly aggressive tumors with CA and associated with worse clinical outcome.** (**A**) Correlation of TACC3 mRNA expression with CA20 score in breast cancer tumors in GSE25066 dataset. (**B**) TACC3 mRNA expression in low vs. high CA tumors in GSE25066 dataset. (**C, D**) Expression of CA20 score (C) and TACC3 mRNA (D) in different breast cancer subtypes in GSE25066 dataset. (**E**) Correlation of TACC3 mRNA expression with CA20 score in cancer cell lines in CCLE dataset. (**F, G**) TACC3 mRNA expression (F) and disease-free survival (G) in low vs. high CA tumors of prostate. (**H, I**) TACC3 mRNA expression (H) and relapse-free survival (I) in low vs. high CA tumors of lung. (**J, K**) TACC3 mRNA expression (J) and overall survival (K) in low vs. high CA tumors of head & neck. CA, centrosome amplification.

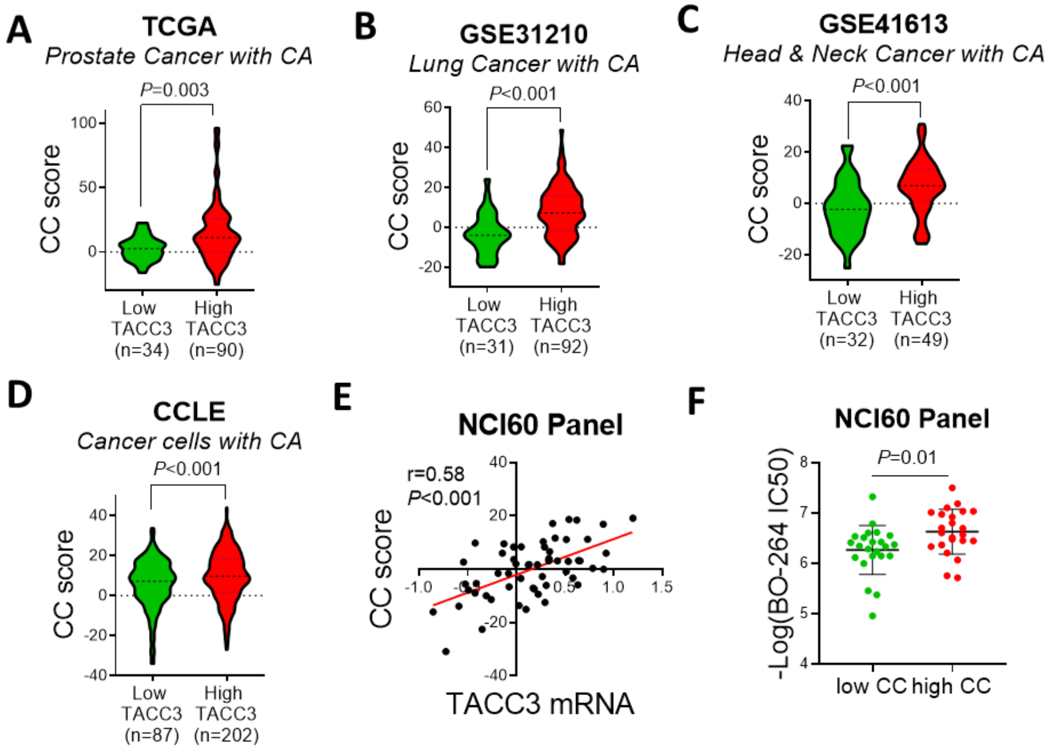


**Fig. S2. TACC3 expression correlates with CC in tumors and cancer cell lines, and high CC correlates with higher sensitivity to TACC3 inhibition.** (**A-C**) Expression of CC score in prostate (A), lung (B), and head & neck (C) tumors bearing CA and separated based on low vs. high TACC3 expression. (**D**) Expression of CC score in cancer cell lines with CA in CCLE dataset separated based on low vs. high TACC3 expression. (**E**) Correlation of TACC3 expression with CC score in NCI60 cell line panel. (**F**) BO-264 IC50 values (-log) in NCI60 cells separated based on CC score.

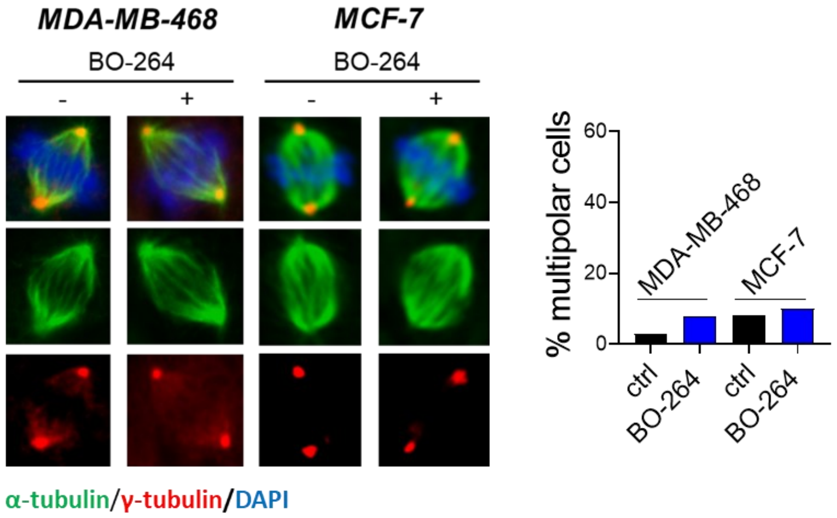


**Fig. S3. TACC3 inhibition-mediated spindle defects are specific to cancer cells with CA and high TACC3 and KIFC1 are associated with worse survival.** Multipolar spindle formation in BO-264-treated MDA-MB-468 and MCF-7 cells as shown by α- (spindle, green) and γ- (centrosome, red) tubulin staining. Quantification of mitotic cells with multipolar spindles is provided on the right panel.

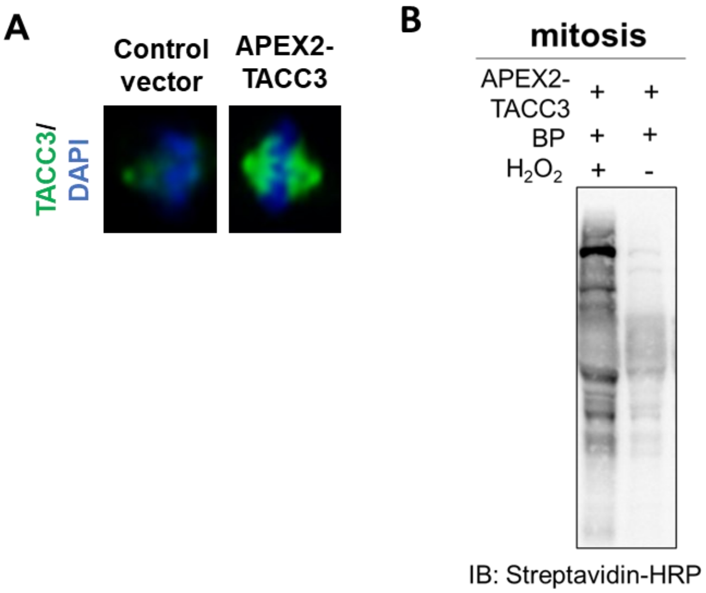


**Fig. S4. Localization of APEX2-TACC3 in mitotic cells and western blot validation of biotinylation.** (**A**) Immunofluorescence staining of APEX2-TACC3 in mitotic JIMT-1 cells. (**B**) Western blot analysis of biotinylated proteins upon $H_2O_2$ in mitotic JIMT-1 cells overexpressing APEX2-TACC3.

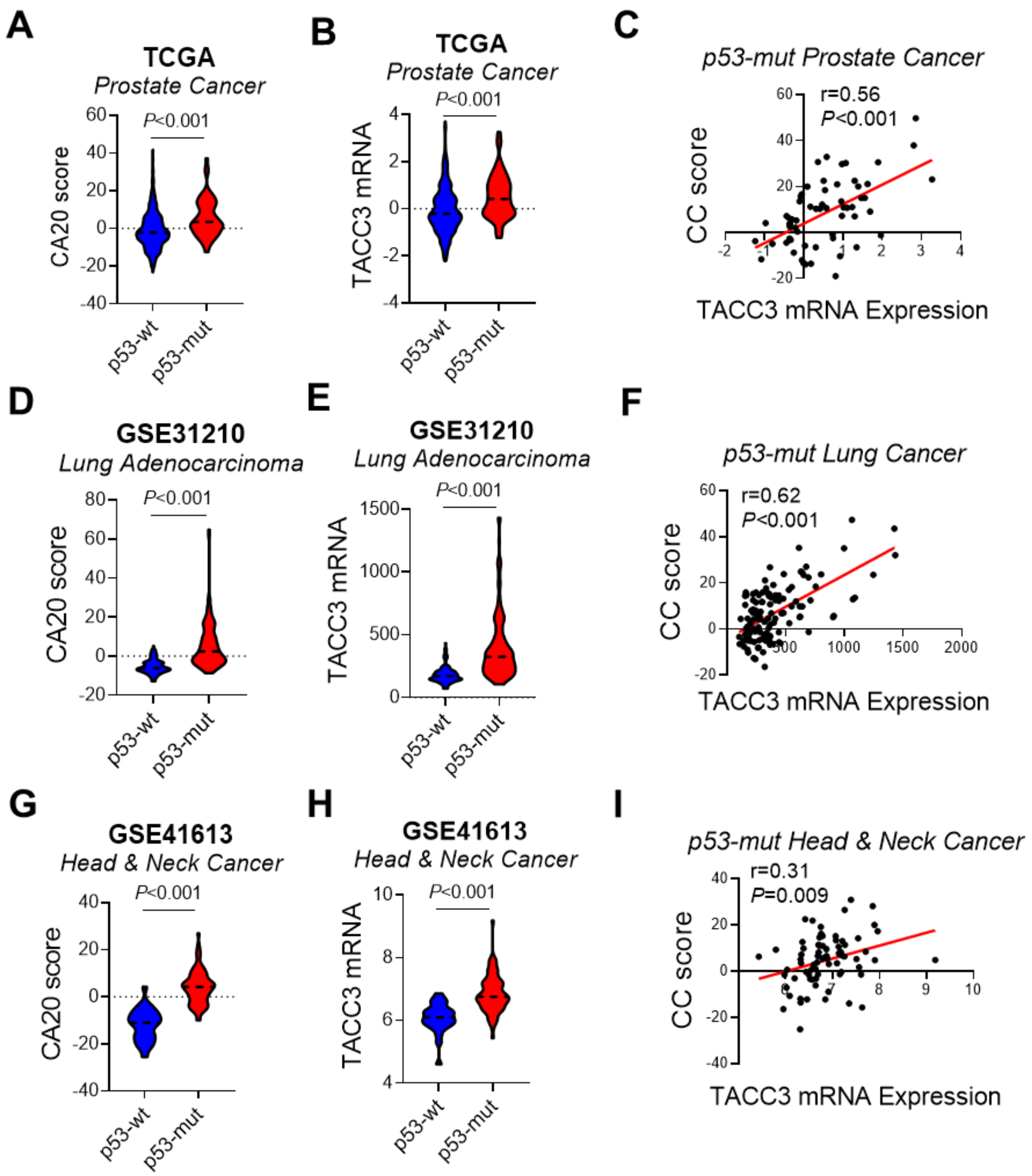


**Fig. S5. Correlation of TACC3 with CC in p53-mut cancer patients.** (**A-C**) CA20 score (A) and TACC3 (B) expression in p53-wt vs. p53-mut prostate cancer patients in TCGA dataset, and the correlation between TACC3 expression and CC score in p53-mut prostate cancer patients (C). (**D-F**) CA20 score (D) and TACC3 (E) expression in p53-wt vs. p53-mut lung cancer patients in GSE31210 dataset, and the correlation between TACC3 expression and CC score in p53-mut lung cancer patients (F). (**G-I**) CA20 score (G) and TACC3 (H) expression in p53-wt vs p53-mut head & neck cancer patients in GSE41613 dataset, and the correlation between TACC3 expression and CC score in p53-mut head & neck cancer patients (I).

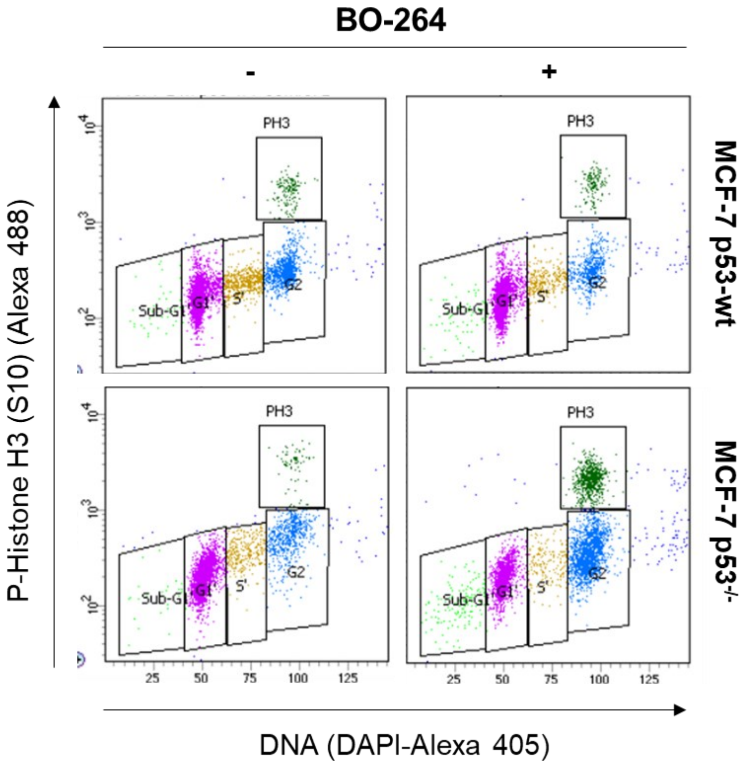


**Fig. S6. Cell cycle analysis to assess the effects of TACC3 inhibition on different cell cycle phases in MCF-7 p53-wt vs. p53$^{-/-}$ cells.** Flow cytometry analysis of DAPI and p-Histone H3 in MCF-7 p53-wt vs. p53$^{-/-}$ cells treated with 2 μM BO-264 for 24 hrs. Population of cells at different cell cycle phases are depicted with different colors and shown in boxes.

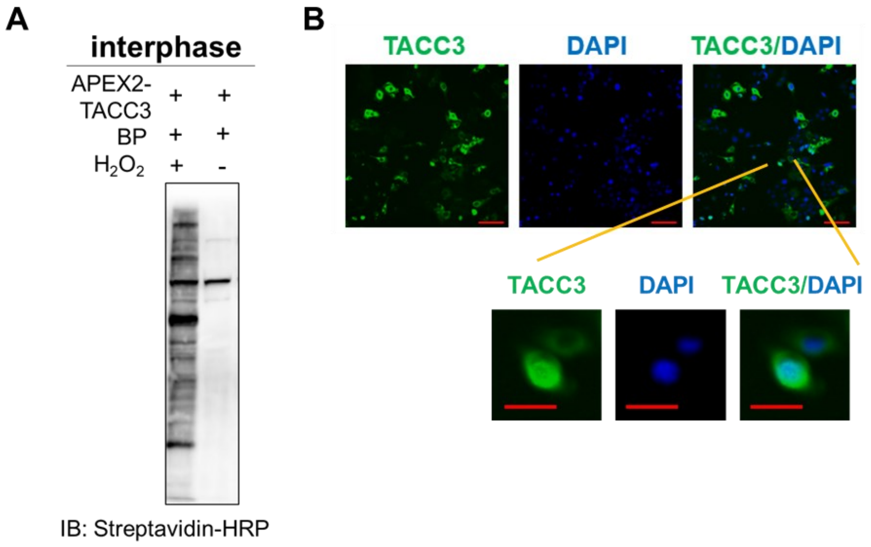


**Fig. S7. Western blot validation of biotinylation and localization of APEX2-TACC3 in interphase cells.** (**A**) Western blot analysis of biotinylated proteins upon $H_2O_2$ in interphase JIMT-1 cells overexpressing APEX2-TACC3. (**B**) Immunofluorescence staining of TACC3 in interphase JIMT-1 cells overexpressing APEX2-TACC3. DAPI is a marker for nucleus.

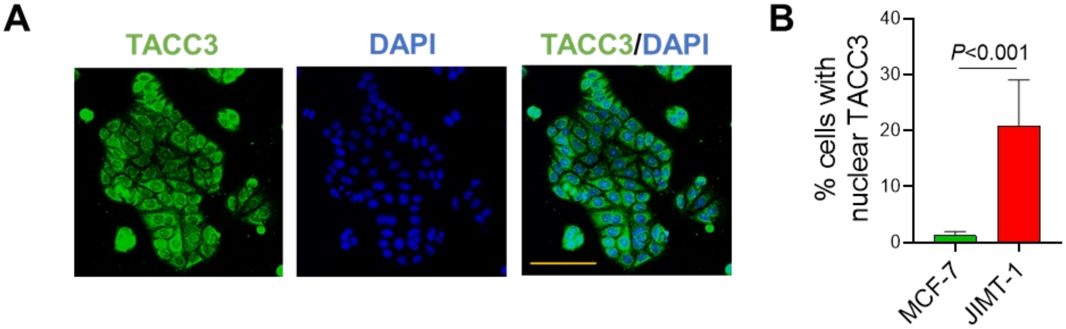


**Fig. S8. TACC3 localization to nucleus is specific to cancer cells with CA.** (**A**) Immunofluorescence staining of TACC3 in MCF-7 cells. DAPI is a marker for nucleus. (**B**) Percentage of non-CA, MCF-7 and CA, JIMT-1 cells with nuclear TACC3 expression.

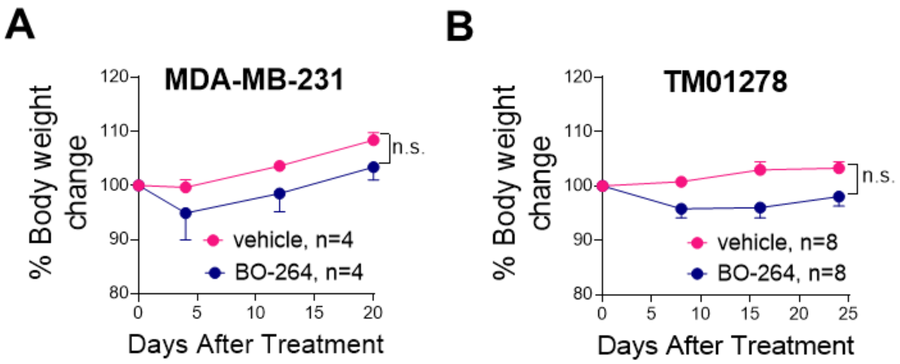


**Fig. S9. Percent body weight change in BO-264-treated MDA-MB-231 xenografts and TM01278 PDXs.** (**A, B**) Percent body weight change of MDA-MB-231 xenografts (A) and TM01278 PDXs (B) upon treatment with BO-264 twice daily at a dose of 75 mg/kg (p.o.). n.s., not significant.

**Table S1.** Sequences of siRNAs used.

| Gene Name | NCBI Gene ID | Company | Catalog number | sequence |
|---|---|---|---|---|
| TACC3 | 10460 | Dharmacon | D-004155-02-0002 | GAGCGGACCUGUAAAACUA |
| KIFC1 | 3833 | Dharmacon | D-004958-02-0002 | GUGCUAAGAUGCUCAUGUU |
| HDAC2 | 3066 | Dharmacon | D-003495-05-0002 | CGGUAUCAUUCCAUAAAUA |
| MBD2 | 8932 | Dharmacon | D-011555-20-0002 | GGGCUAAGUGCUGGCAAGA |

**Table S2.** Sequences of qRT-PCR primers.

| Gene Name | NCBI Gene ID | | Primer sequence |
|---|---|---|---|
| ACTB | 60 | Forward | 5'-CCAACCGCGAGAAGATGA-3' |
| | | Reverse | 5'-CCAGAGGCGTACAGGGATAG-3' |
| HPRT | 3251 | Forward | 5'-TGACCTTGATTTATTTTGCATACC-3' |
| | | Reverse | 5'-CGAGCAAGACGTTCAGTCCT-3' |
| DAPK1 | 1612 | Forward | 5'-TGTCTTCCACCAACTCCAGCAG-3' |
| | | Reverse | 5'-AAATCGCCAACTCCATTCAAATAAGC-3' |
| KLK10 | 5655 | Forward | 5'-GCCCGGAGAGTGAAGTACAA-3' |
| | | Reverse | 5'-GTAAACACCCCACGAGAGGA-3' |
| APAF1 | 317 | Forward | 5'-CACGTTCAAAGGTGGCTGAT-3' |
| | | Reverse | 5'-TGGTCAACTGCAAGGACCAT-3' |
| CDKN1A | 1026 | Forward | 5'-TGAGCCGCGACTGTGATG-3' |
| | | Reverse | 5'-GTCTCGGTGACAAAGTCGAAGTT-3' |
| CDKN2A | 1029 | Forward | 5'-GAGCAGCATGGAGCCTTC-3' |
| | | Reverse | 5'-CCTCCGACCGTAACTATTCG-3' |

**Table S3.** List of antibodies used in Western blot (WB), immunofluorescence (IF) and immunoprecipitation (IP) experiments.

| Antibody | Provider | Catalog number | WB dilution | IF dilution | IP dilution |
|---|---|---|---|---|---|
| Alexa Fluor® 488 anti-mouse | Life Technologies | A-11001 | - | 1:200 | - |
| Alexa Fluor® 647 anti-rabbit | Life Technologies | A-31573 | - | 1:200 | - |
| Beta-actin | MP Biomedicals | 691001 | 1:10000 | - | - |
| Cleaved PARP | Cell Signaling Technology | 5625 | 1:1000 | - | - |
| Alpha-tubulin | Santa Cruz | 32293 | - | 1:500 | - |
| Gamma-tubulin | Sigma Aldrich | T3195 | - | 1:200 | - |
| p-Histone H3 (Ser10) | Cell Signaling Technology | 4056 | 1:1000 | - | - |
| TACC3 | Santa Cruz | 376883 | 1:1000 | 1:400 | 1:10 |
| KIFC1 | Abcam | ab172620 | 1:1000 | 1:400 | 1:100 |
| MBD2 | Abcam | ab188474 | 1:1000 | 1:400 | - |

| | | | | | |
|---|---|---|---|---|---|
| HDAC2 | Abcam | ab16032 | 1:1000 | 1:400 | - |
| p-Bcl2 | Cell Signaling | 2827T | 1:1000 | - | - |
| Bcl2 | Cell Signaling | 15071T | 1:1000 | - | - |
| Clathrin | Abcam | ab21679 | 1:1000 | - | - |
| GFP | Genetex | GTX628528 | 1:1000 | - | - |
